# Causality-aware graph neural networks for functional stratification and phenotype prediction at scale

**DOI:** 10.1101/2024.12.28.630070

**Authors:** Charalampos P. Triantafyllidis, Ricardo Aguas

## Abstract

We employ a computational framework that integrates mathematical programming and graph neural networks to elucidate functional phenotypic heterogeneity in disease by classifying entire pathways under various conditions of interest. Our approach combines two distinct, yet seamlessly integrated, modeling schemes. First, we leverage Prior Knowledge Networks (PKNs) to reconstruct gene networks from genomic and transcriptomic data. We demonstrate how this can be achieved through mathematical programming optimization and provide examples using comprehensive established databases. We then tailor Graph Neural Networks (GNNs) to classify each network as a single data point at *graph-level*, using various node embeddings and edge attributes. These networks may vary in their biological or molecular annotations, which serves as a labeling scheme for their supervised classification. We apply the framework to the human DNA damage and repair pathway using the *TP53* regulon in a pancancer study across cell-lines and tumor samples to classify Gene Regulatory Networks (GRNs) across different *TP53* mutation types. This approach allows us to identify mutations with distinguishable functional profiles which can be related to specific phenotypes, thus providing a data-driven pipeline for genotype-to-phenotype translation. This scalable approach enables the classification of diverse conditions within the multi-factorial nature of diseases and disentangles their polygenic complexity by revealing new functional patterns through a causal representation.

## Introduction

The sparsity of causal interpretation in medical sciences [1, 2, 3] and the need to utilize it using high-throughput genomic and transcriptomic data combined with the wider availability of computational resources and modeling approaches, marks a distinct gap in our understanding of disease [4, 5, 6, 7]. To address this gap, particularly with regard to the biological deregulation of genomic underpinnings, it is instrumental to elucidate how signals propagate across molecules from root(s) to target(s). Genes and the proteins they encode can be effectively captured in a computational model by an artificial *wiring* system, resulting in network representations or *graphs* [8, 9, 10, 11, 12, 13]. These mathematical objects contain nodes (genes in this context) and edges (directed interactions typically mapping an activating or inhibitory effect) indicating the Mode of Regulation (MoR). Genes can interact in a bi-directional way but also in synergy to express a particular phenotype, a cellular process carrying out specific biological tasks (such as DNA damage and repair, proliferation, apoptosis and others). The importance of this mathematical *language* to communicate gene interactions is rooted in its ability to provide causal interpretation of the underlying shared effects [14] by using directed acyclic graphs (DAG).

Thus, a systems biology approach using high-throughput data and advanced computational methods is instrumental in revealing the functional implications of different conditions that can map genotypes and transcriptomic data to phenotypes in disease. In particular, graph-based models provide a natural representation of gene regulatory networks [15, 16], enabling the integration of multi-dimensional data to capture structural and therefore functional differences across different layers such as gene activity profiles and regulatory direction and modes.

This line of work builds on two primary blocks, incorporating distinct and yet seamlessly integrated modeling schemes. The first one is based on well-established Mixed-Integer Linear Programming (MILP) models [17] which we employ to map expression data to PKNs and hence reconstruct their topology and gene activity profiles for the pathway of interest. These models are deterministic and global optimal, offering advantages in the optimization and reconstruction of the networks, in terms of computational efficiency, scaling and stability. Furthermore, a plethora of network analysis metrics such as community detection and centrality measures, can be applied to the reconstructed networks to provide further insights of possible biological and functional implications. When we incorporate Prior Knowledge Networks (PKNs) derived from established databases or literature-curated sources, a common practice is to associate each interaction with a confidence score, reflecting the strength or reliability and possible distance from ground-truth. In the context of transcription factors (TFs) and their regulon, such confidence scores represent the degree of experimental or computational validation that supports the regulatory relationship between a TF and its putative target gene(s). Starting from an unpruned base PKN, consisting of as many target genes as possible, we then seek to reduce the number of edges/nodes in our network topology by minimizing the mismatch between that prior knowledge and transcriptomic data for a targeted regulon (a pancancer regulon in our case) for each sample. The resulting pruned networks provide the input to classify the pathway represented at graph-level, by using Graph Neural Networks (GNNs).

This study builds on the computational innovations of different modeling approaches to explore in specific the regulatory impact of somatic mutations in *TP53* as a paradigm, and to extend the functional heterogeneity of pathways of interest across different conditions. It differs from previous results [18] in many respects. Firstly, the original analysis used undirected graph edge intersection comparison to assess differences or similarities as a first measure of comparison across same conditions (e.g. mutation types). Secondly, the current study is the first to also incorporate the activity of the nodes (the gene activity profiles) across the graphs, not only comparing topologies as in [18], but also the edge type as mode of regulation (inhibition, activation). Additionally, the feature engineering for the GNN classifier implemented here is more detailed, able to capture high level structural and biological expressiveness in the networks, given not only the mechanics behind message passing in GNNs but also the ability to highlight certain genes of interest (see Methods for the spotlight mechanism). This additional layering of various node and edge attributes calculated on a graph-level can further identify key graph attributes but also subtle differences emerging from the biological heterogeneity that may be imposed across the conditions of interest analyzed. Finally, the community detection method used in this line of work accepts directed graphs as opposed to the methods used in the previous study, where the input graph had to be converted to undirected first, eliminating direction of regulation and therefore naturally causing loss of biological knowledge.

By reconstructing and optimizing GRNs from the Cancer Cell Line Encyclopedia (CCLE) [19] dataset and The Cancer Genome Atlas (TCGA) [20], we aim to elucidate mutation-driven differences in regulatory architecture. The integration of graph-based classification and network-level analyses allows us to investigate whether somatic mutations alter regulatory networks to a detectable degree, by finding a classification boundary. The combination of these two modeling schemes, the network reconstruction and the GNN classifier, yields the integrated framework illustrated by Figure 1. To demonstrate the applicability and merit of the approach, we apply it to a multi-factorial disease, cancer, particularly due to the entwined causality of its genomic nature. The insights gained not only advance our understanding of transcription factor biology but also hold potential for identifying novel therapeutic targets and biomarkers entwined to precision oncology, and expand further outside of our case study to multiple diseases, given the availability of data.

**Figure 1:**
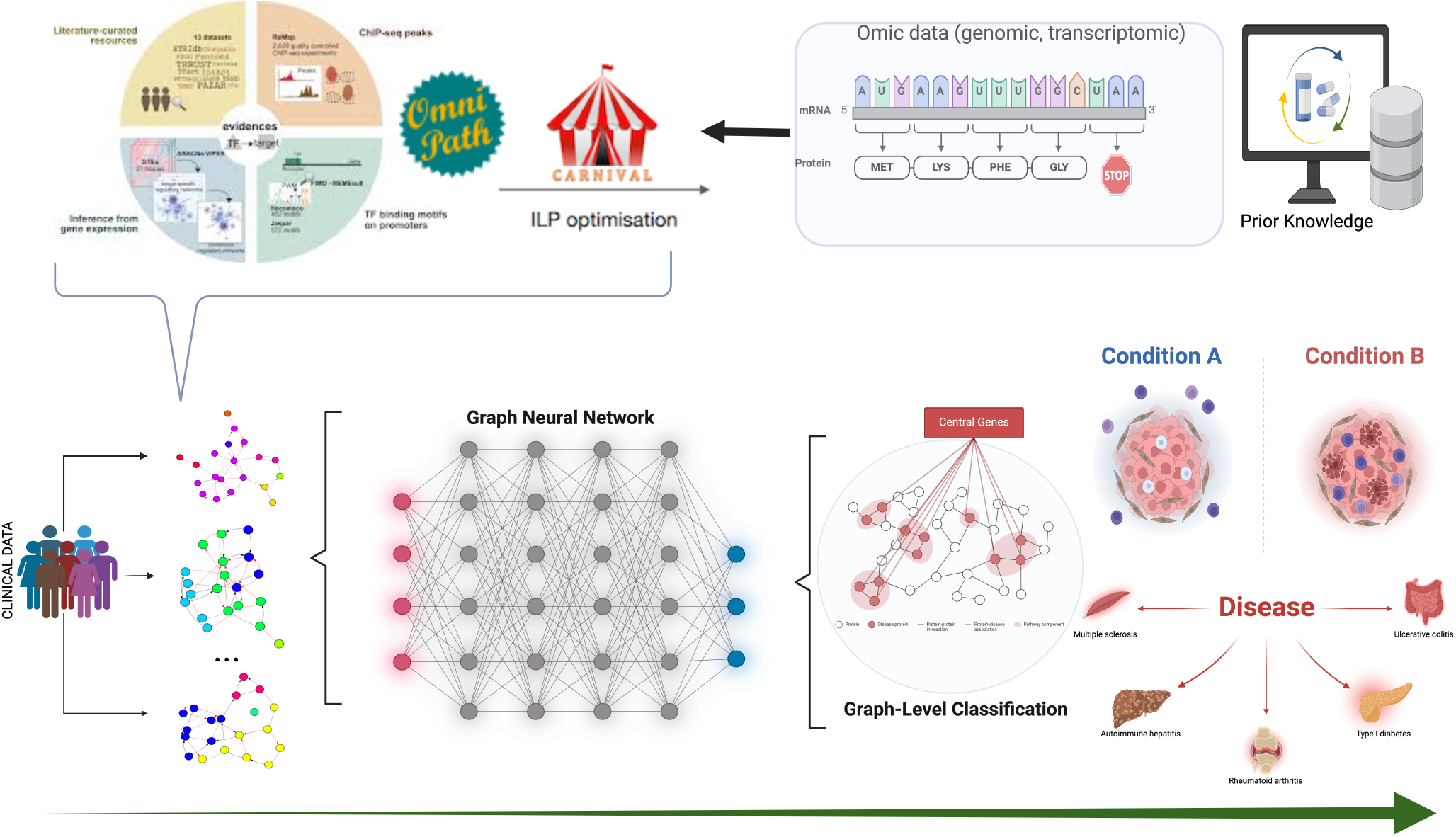
The developed framework workflow. Prior Knowledge Networks (PKNs) and omics data are combined to provide the input for optimization models to reconstruct gene regulatory networks across pathways of interest, in this case the *TP53* pathway and its regulon. These optimized networks are then annotated (labeled) across conditions of interest as they are reconstructed per sample; in this example, mutation types of the transcription factor *TP53*. We then classify the networks across these labels using Graph Neural Networks that can distinguish entire pathways as a single data point (graph-level classification) to predict disease, or states of diseases (onset, progression, severity, etc.), or any other event of interest. Created in ©bioRender.

In conclusion, we provide a computational approach to causal inference in the endeavor to identify phenotypic diversity directly linked to disease states (for example mutations, severity, onset, progression, metastasis for cancer etc.) and demonstrate how we can disentangle the complex molecular and biological underpinnings of multi-factorial diseases using networks and machine learning/optimization. An application in a pancancer study as a demonstrating paradigm shows the applicability and utility of the framework as a first step in using the proposed pipeline. To contextualize our framework within existing approaches, we next review recent efforts applying deep learning models, particularly GNNs, in genomic and pathway analysis.

### Related work

Deep learning models have become a key approach in analyzing graph-structured data, enabling the extraction of meaningful patterns from complex networks of various types and origin. Specifically for cancer and *TP53*, multiple deep learning models have been deployed. For example, in [21] deep learning has been successfully used to identify resistance to therapy p53-pathway mediated mechanisms and related combinatorial targets for lung cancer. Additionally, in [22], a Recurrent Neural Network (RNN) has been employed to illuminate p53 temporal interaction dynamics. Furthermore, a GNN approach has been utilized in [23] to predict protein stability change given novel gene mutations. Gene module dissection using GNNs has been also recently demonstrated in [24] and gene signatures have also been extracted using Graph Convolution Networks specifically for *TP53* mutations in [25], or to predict missense driver mutations [26].

Graph Neural Networks [27, 28, 29, 30, 31, 32] represent a cutting-edge special class of deep learning models, specifically developed for graph-based structured data, where other renditions of neural networks cannot be applied. This design feature renders them ideal for interrogating GRNs. GNNs provide a powerful framework for modeling complex relational structures inherent in a biological setting. Traditional machine learning approaches often overlook the rich dependencies between biological entities such as genes and proteins, treating data as independent. GNNs address this limitation by explicitly leveraging graph structures to learn from these relationships, enabling a range of predictive and interpretative tasks across healthcare and biomedical research, including network reconstruction [33], node prediction and classification [34, 35], link prediction [36], disease diagnosis/prediction [37, 38] and graph-level classification [39].

In genomics, GNNs are increasingly applied in drug–target prediction [40], modelling cell to cell interactions [41], and predicting functional roles of genes within biological systems [42]. In clinical settings, constructing patient similarity graphs from electronic health records (EHRs) enables GNNs to predict disease risk, treatment responses, and recommend medications [43, 44]. Recent advances also demonstrate the utility of GNNs in the extraction of novel pairs of extracellular interacting genes using spatial transcriptomics [45].

More specifically, relevant to this work, the task of Graph-level classification using Graph Neural Networks (GNNs) has gained significant attention. Entire biological networks, such as signaling pathways, metabolic circuits, or patient-specific GRNs, are treated as single graph objects to be classified based on phenotypic or clinical labels. This approach enables applications like distinguishing disease-specific pathway topologies, identifying disrupted regulatory circuits in cancer, and predicting clinical outcomes from patient-derived GRNs. For instance, PathGNN captures topological features of pathways to predict long-term survival in various cancers [46]. Similarly, GNN4DM identifies overlapping functional disease modules by modeling disease-associated gene networks [47]. Directed Graph Convolutional Networks have been proposed for GRN inference, effectively handling directed graph structures inherent in regulatory networks [48]. Additionally, multi-level GNN frameworks integrate gene-level and pathway-level information to stratify tumor risk, leveraging multi-omics data for comprehensive analysis [49]. These methodologies underscore the versatility of GNNs in capturing both molecular activity and network organization, advancing personalized medicine and biomarker discovery.

The methodology presented here shares similarities but also key differences with previous related studies. We tailor GNNs by employing a GATv2Conv model, which effectively captures directed graphs as inputs, but simultaneously allows for the incorporation of edge attributes, which in this case are the modes of regulation. The construction of the input graphs is based on mathematical programming, ensuring we can effectively map transcriptomic data and PKNs to newly reconstructed graphs with altered topology to fit the per-sample mRNA expression each time, instead of the standard GRN approach. The node activities and various features have been engineered to allow the embedding of multiple biologically relevant features, but also features engineered to assist dissecting minor and major topological differences across the input graphs, for example community detection methods. Gene activity profiles have not been utilized in previous studies to assist the graph-level classification. Additionally, a further spotlight mechanism has been introduced here tailored to attend to genes of interest, emphasizing in single gene dominant regulation or complex antagonistic gene sets behavior. All these modifications result in significant improvement compared with previous approaches, mainly highlighting not only various low level technical considerations but also the focal point of the proposed framework, which is to simulate entire pathways under a condition of interest as a single data point to be classified by using a GNN. Having established the computational context, we now focus on the biological background of our case study, *TP53* in cancer.

### Cancer and the *TP53* regulon

Cancer onset and its rapid progression is a result of pervasive biological deregulation taking place across a plethora of cellular processes [50, 51, 52, 53]. As aberrant gene expression drives oncogenic processes and initiates a multitude of neoplastic diseases [54], the master regulators of the genome, known as Transcription Factors (TF) [55, 56, 57], dictate the gene activity profile of many downstream targets, known as their *regulon*. Knowledge of the intricate dynamics of the known 1,600 DNA-binding TFs [58], is paramount in expanding our understanding of gene expression programs, transcription and post-transcriptional regulation in disease. Their perturbations in cancer emanating from stress conditions such as mutations, irradiation, hypoxia [59, 60] or drug effects and other stimuli, are pivotal in determining a cascade of processes underlying disease onset, severity and progression.

Among these transcription factors, *TP53* [61, 62, 63] stands out due to its widespread impact on multiple types of cancer [64, 65], influencing crucial pathways for cell cycle control, apoptosis and DNA repair, and hence aptly named the *Guardian of the Genome* [66]. Somatic mutations in TFs [67] can alter the functional form of proteins and their ability to dock and perform cellular tasks, leading to abnormal states. In *TP53*, they manifest as heterogeneous effects on downstream gene expression [68, 69] and can be used as annotated conditions of interest in the developed framework. Besides mutations, any external stimuli that can be described as a perturbing effect could also be considered, such as hypoxia and irradiation, as demonstrated in [18], or for example different drug concentrations/effects. *TP53* as a major transcription factor involved in cellular defenses against cancer is inactivated in nearly all tumors, with mutations in *TP53* occurring in approximately 50% of human cancers [70]. Tumor suppressors like *TP53* typically experience loss-of-function mutations. Unlike other tumor suppressors, *TP53* mutations are predominantly missense variants concentrated in the DNA-binding domain (DBD). *TP53*’s diverse mutation landscape is a result of its structural fragility, evolutionary compromises, and intrinsic mutability [70, 71]. Most mutations lead to loss of tumor-suppressive function, with additional dominant-negative or aggregation-related effects exacerbating tumorigenesis. While some mutations exhibit neomorphic effects, these are often secondary to the primary loss of p53 function [72].

This functional heterogeneity poses a significant challenge for understanding mutation-specific effects, requiring the dissection of the molecular mechanisms driving distinct regulatory states. Analysis of the functional implications in *TP53* state support that Wild-Type (WT) status enhances both innate and adaptive immunity by promoting pro-inflammatory cytokine secretion, activating the cGAS-STING pathway and regulating antigen presentation via MHC-I, influencing immune checkpoint pathways [73]. Additionally, *TP53* mutations can create an immunosuppressive tumor micro-environment (TME) with altered cytokine and chemokine production that recruits immunosuppressive cells like myeloid-derived suppressor cells (MDSCs) and M2 macrophages and induction of chronic inflammation via NF-*κ*B and other signaling pathways, allowing tumor progression [73]. Further investigation of *TP53* mutations reveals their impact on tumor suppressing ability, and shows that [74] missense mutations are associated with destabilization of p53’ s DNA-binding domain (DBD) and oligomerization domain (OD). This offers potential therapeutic targets, and suggests that p53 function could be recovered by restoring DBD and OD stability by other means such as drug delivery. Specific missense *TP53* mutations can be predicted with extremely high accuracy [75], suggesting there is significant and identifiable heterogeneity across mutations each of which in turn contributes differently to p53 functional activity. Finally, *hotspot* mutations, mutations of a specific protein change with very high occurrence, were strongly associated with loss of function. The complex impact of *TP53* somatic mutations delineates an ideal application landscape for the purposes of the proposed computational framework.

## Results

We first explored the heterogeneity found in the expression profile of the pancancer regulon to ensure consistency with the finding in [18]. We extracted all cell line samples carrying a single *TP53* mutation and combined the expression levels of its regulon with the mutational profiles and other molecular information such as cancer type, protein change, hotspot and deleterious function of the mutation for each sample. We then stratified the expression levels of the regulon per mutation type and performed a dual hierarchical clustering of Z-standardized transformed expression levels to understand whether either cell-lines, or gene targets of *TP53* cluster together in a consistent pattern, Figure 2. No pattern was observed across either WT or mutated samples, emphasizing the transcriptional divergence of *TP53* with the potential functional implications remaining unresolved by isolated expression analysis. The lack of discernible patterns in raw expression motivated the use of reconstructed regulatory networks, which incorporate both expression and prior knowledge about the regulatory networks to potentially reveal functional diversity. Transcription factor activation is known to be observable only in the gene activity profile of the regulon and not in its own expression levels.

**Figure 2:**
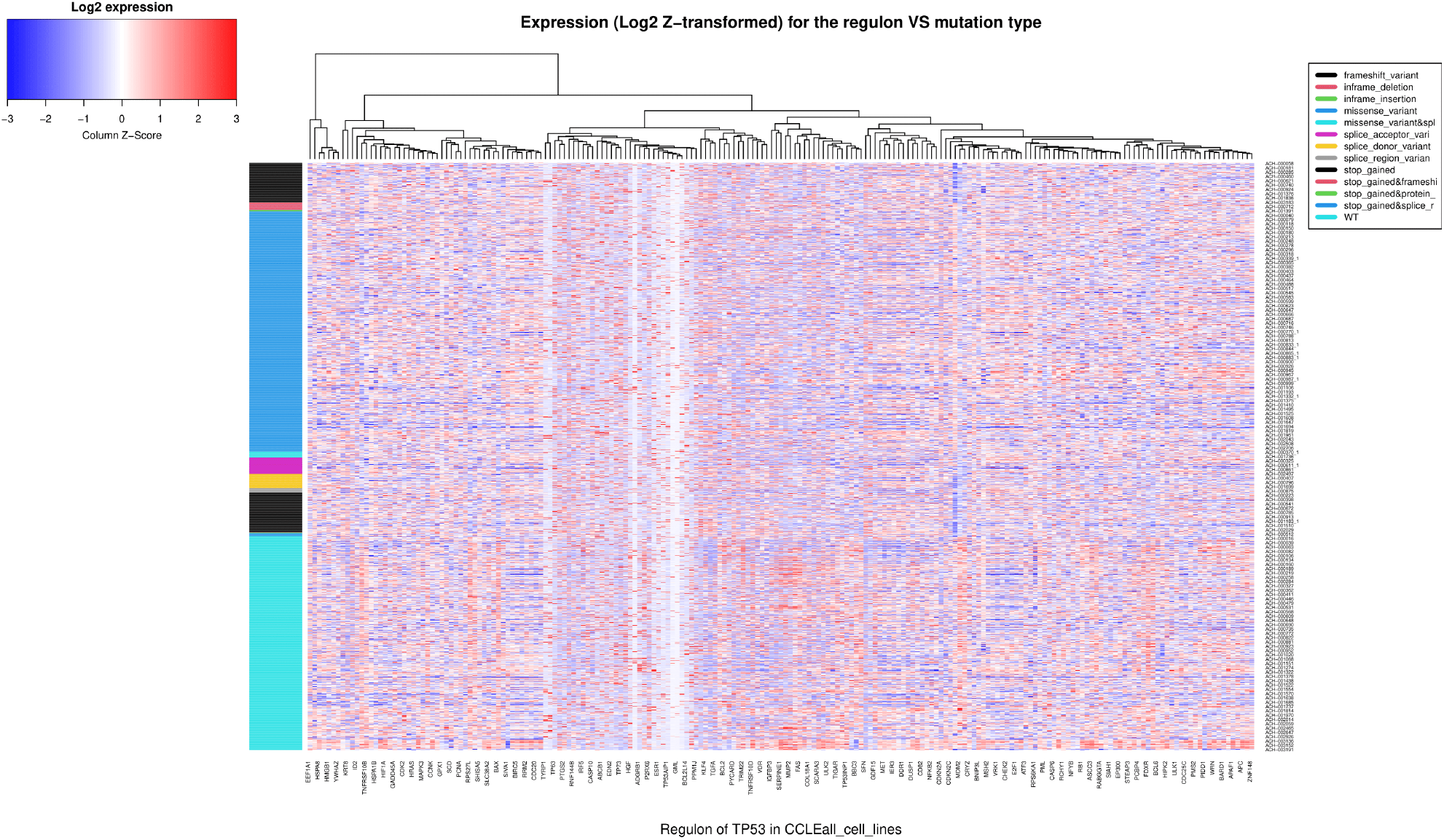
Z-normalized levels of expression for the regulon (set of target genes) of transcription factor *TP53*, as columns are depicted in this heatmap plot and samples (i.e. unique mutation of *TP53* per sample as rows. We stratify the samples/rows in the *TP53* mutation type using a colored legend. We can see that missense mutations are prevalent, but WT samples are also dominant. Approximately 1600 samples are included in this figure, pancancer from CCLE 24Q2.

Distributions of mutation types for *TP53* across the datasets are given in Figure 3-(A-B). Missense mutations and WT samples dominate occurrence across both cell lines and tumor samples. We utilized the deleterious function of *TP53* mutations to model its perturbing effect in the network each time. We consider as deleterious the effect of rendering the gene (subsequently its protein) dysfunctional, and so, a deleterious mutation would implicitly mean that *TP53* is deactivated. This perturbing effect is used as an input in the CARNIVAL pipeline from which we obtain insights into the classification and expression patterns of *TP53* related mutations across various cell lines. For CCLE e.g., Figure 3-C shows the expression levels of *TP53* across mutant (and stratified across deleterious function of mutation) and WT models (cell-lines). Statistically significant (*p* < 2.2 *e* −16, Wilcoxon and T-tests) higher expression is observed across deleterious and mutant models versus non-deleterious mutation cell-lines. WT samples exhibit similar expression patterns to the high mutant deleterious models, again confirming that expression levels for *TP53* transcription factors are not representative of status (active/inactivated). Figure 3-D shows the violin plot of *TP53* expression stratified by mutation type and WT samples included in CCLE, and significant differences across the means of the variants.

**Figure 3:**
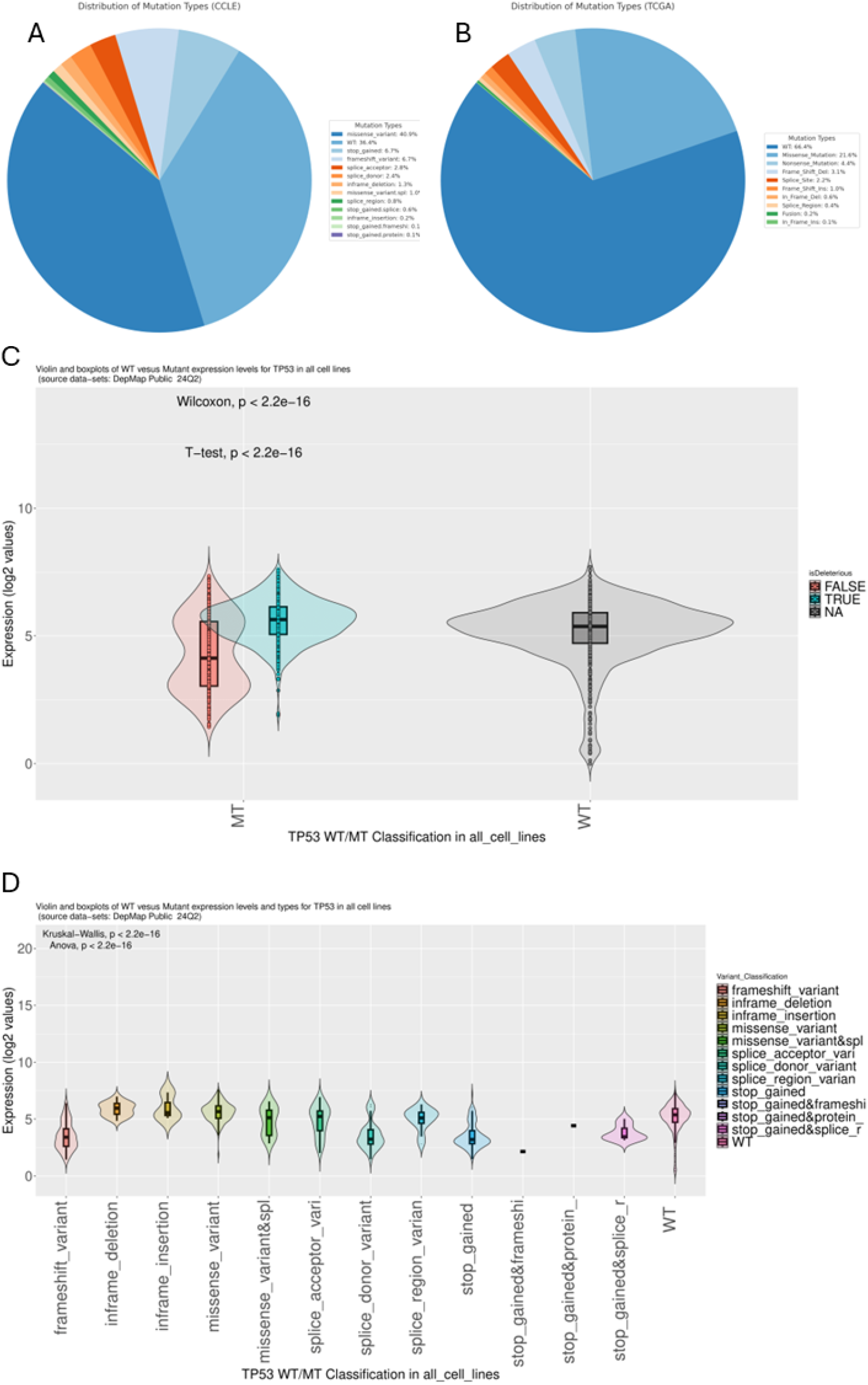
**(A-B)** Distribution of mutation types across the datasets used. Although WT and missense mutations dominate both cases, a slight variation is observed across the datasets, where in tumor samples WT samples are more prevalent than in cell lines. In CCLE, almost 40% of the samples carry a missense TP53 mutation, and 36.4% of the samples are *T P* 53^*WT*^. In TCGA, almost 66% of the samples are *T P* 53^*WT*^, and from the rest 34%, more than half (21.6%) carry a missense TP53 mutation. **C)** Deleterious function of mutation stratified across mutated samples and contrasted with controls (WT). We observe a statistically significant difference in the means of log2 normalized expression for *TP53* for the deleterious mutations against non-deleterious, and higher than WT expression. **D)** A violin and box-plot analysis compares the expression levels of the *TP53* gene across different mutat7ion types in various cell lines.The Kruskal-Wallis and ANOVA tests indicate a highly significant difference in expression values across groups (*p* < 2.2 *e* − 16).

### Network analysis of reconstructed networks reveals functional differences

The initial backbone PKN, as extracted from *OmniPathR*, contains approximately forty thousand interactions. This graph serves as the building block for every CCLE or TCGA model being re-optimized as a single gene regulatory network, using expression of each single sample as input. In essence, we reconstruct as many gene regulatory networks as cell-lines, or CCLE/TCGA samples, that contain data available to characterize *TP53* status, both in terms of expression and also the deleterious function of the mutation, each time imposing the corresponding perturbing effect.

The initial PKNs including all downstream identified signed and directed interactions with *TP53*, have been reconstructed using the full expression matrix and *TP53* perturbation from the deleterious function of mutation each time (deleterious:inactive). For WT samples we assume *TP53* is both functional and active. We used CARNIVAL to optimally reconstruct the networks, which then served as an input of training graphs for the GNN classifier. The network analysis we performed included three parts: i) basic plots to analyze the networks across an array of different features, ii) community detection to identify densely connected hubs, and iii) betweenness centrality analysis per identified hub and extraction of unified signatures across each mutation type (See Methods) with the inclusion of gene status (active/inactive). The violin plots (Suppl. Figures S1,S2) indicate that for both CCLE and TCGA, and across the different metrics tested, missense mutations and WT samples account for the majority of the detectable variability. Figure 4 shows a hierarchical dual clustering with the regulon target genes on the x-axis and the mutation types plus WT samples as different categories on the y-axis, for the CCLE dataset. The genes can be absent (white), active (blue) and inactive (red), across each mutation type / WT sample. We can observe high dissimilarity between the majority of the extracted signatures. These signatures highlight functional differences across the various mutation types, resulting in structural differences of the reconstructed GRNs.

**Figure 4:**
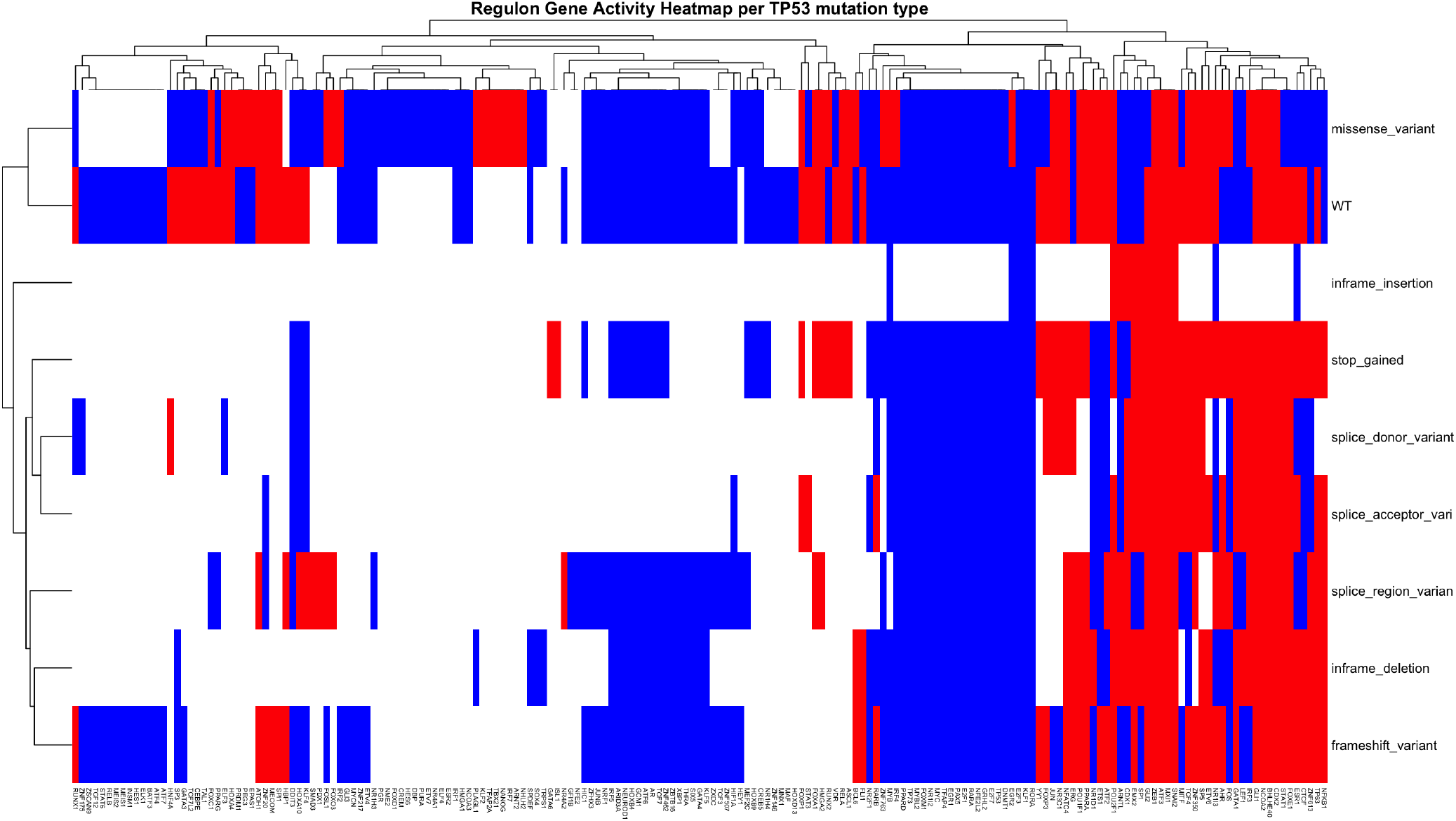
Plotting the gene signatures extracted from the community detection analysis in CCLE across each mutation type and WT, and showing gene activity each time. All networks carrying the same *TP53* mutation were stratified into groups. Community detection was performed for each group, and a centrality (maximum) score was used to identify one representative gene per community. The overlap of these sets across the same mutation type or WT each time, is the set shown for each mutation type/WT here respectively. We observe functional network similarity between WT and missense mutations, which may reflect recent findings in the functional landscape of *TP53* mutational variability, where many missense mutations phenotypically behaved as WT-like [76]. The heatmap shows clustered gene status, with genes on the y-axis and samples (or conditions) on the x-axis. Blue indicates up-regulated genes, red denotes down-regulated genes, and white represents absence.

### GNN *learns* graph-level GRNs based on condition of interest (mutation type)

We trained our GNN classifier across two datasets (CCLE and TCGA), using 70% of the samples for training and the remaining for testing. The input graphs contain a wealth of node and edge attributes which we utilized to engineer low and high level features (See Methods). Across those node embeddings, we also incorporated the edge types per mode of regulation each time (inhibition, activation). Our pipeline reads the input graphs and then dissects the node and edge features to prepare the dataset for training and testing. The feature matrix is passed to the graph neural network which utilizes a GATv2Conv model. This rendition allows for seamless integration with directed graphs and also allows for the edge attributes vector to be defined by the user, thus making it ideal for our purposes. The GNN model contains multiple hidden layers and a mini GATv2Conv specifically to focus on genes of interest given by the user. For example, as *TP53* plays a key role in our underlying reconstructed GRN graphs, one might want to highlight the close, hop-dictated, neighborhood of the gene to enable the classifier to provide additional attention to it. This *spotlight* mechanism has been used to provide robust biological underpinnings to our training process. Our heterogeneous graphs were then used to build the classifier by minimizing the cross-entropy function as the task of graph classification imposes.

The TCGA model achieved a misclassification rate of ∼ 16%, while the CCLE model achieved *∼*31%, at best, given our hyper-parameter tuning. The GOI spotlight did not alter the classification efficacy in CCLE, however it introduced a clear 16% gain in classification (16% missclassification, down from 32%) in TCGA when we included two genes of interest, namely *TP53* and *MYC*.

The confusion matrices for each dataset (Suppl. Figure S3) reveal the classification performance for different mutation types across the datasets, showing strong accuracy in predicting WT (wild-type) and missense classes, indicated by the high diagonal values. However, notable misclassification occurs for other less prevalent types of mutations even with balanced label split in the test and train sets, and weight balancing in the loss function, suggesting either a lack of strong signal to differentiate these graphs from others, or the result of small training examples to promote efficient learning. Suppl. Figure S4 shows the Receiver Operating Characteristic (ROC) curve for the multi-label classifier across all variants and WT samples in each of the datasets. Generally, the classifier is able to differentiate across samples with strong presence in the training data, whilst struggling to separate cases low in incidence. In both CCLE and TCGA, the vast majority of samples belong to either missense or WT samples. Scatter plots were generated for the true and predicted class (Figure 5) for both datasets, and the classification accuracy (blue:correct, red:misclassified). Figure 6, shows an interactive plot generated by *HarmonizeR*, our implemented platform in *R*. Here, each data point displays the optimized network (a single sample of CCLE or TCGA) showing both the directionality of interactions and the activity profile (indicating gene activity status, with blue denoting activation and red denoting inactivity).

**Figure 5:**
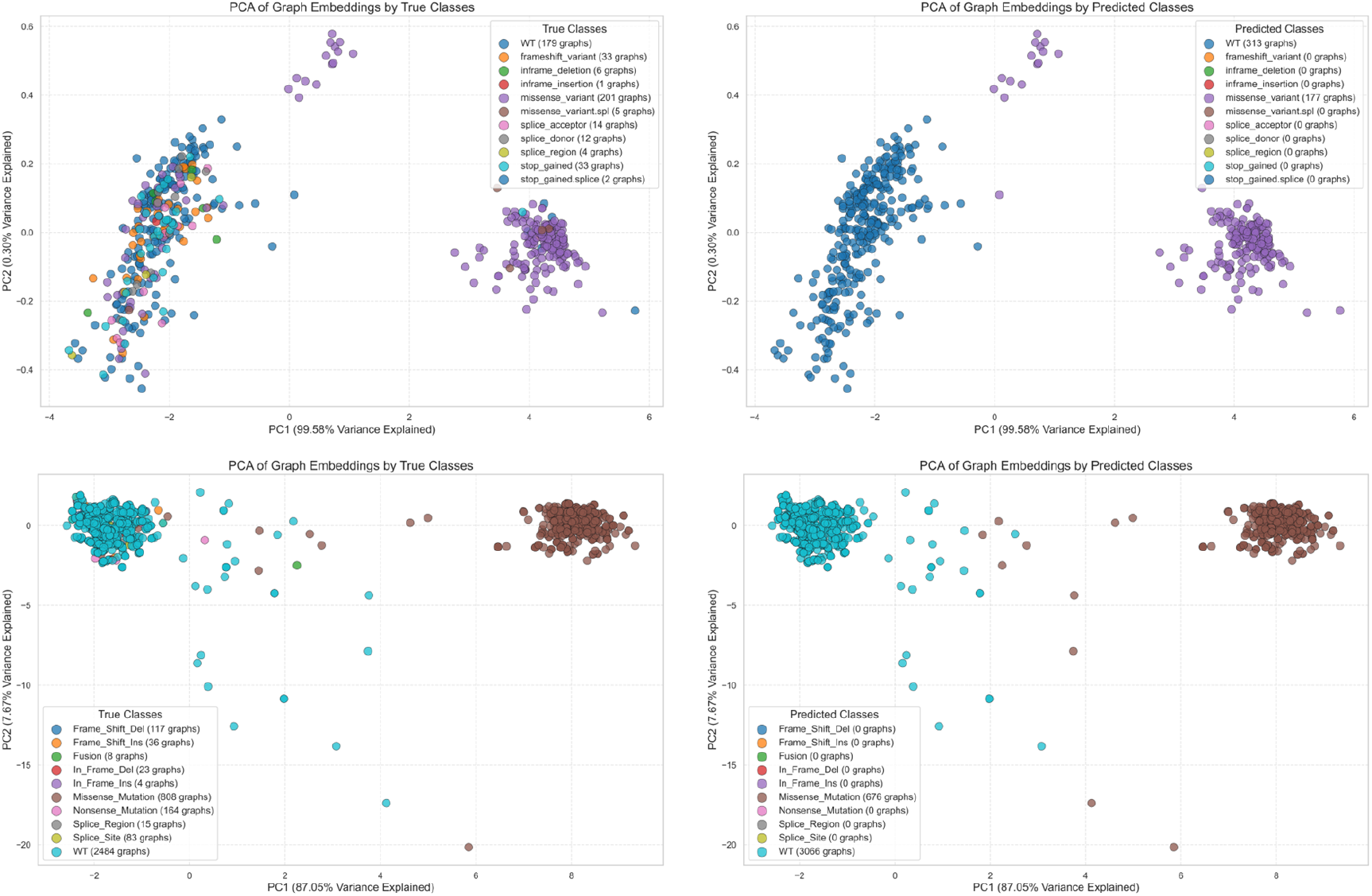
PCA 2-D scatter plots of the learned embeddings with different color annotation for the classified labels, correspondingly. CCLE (top row) and TCGA (bottom row). The right columns show the distribution of the embeddings from the graph neural network output as predicted classes and the left columns show the true labels of the graphs.

**Figure 6:**
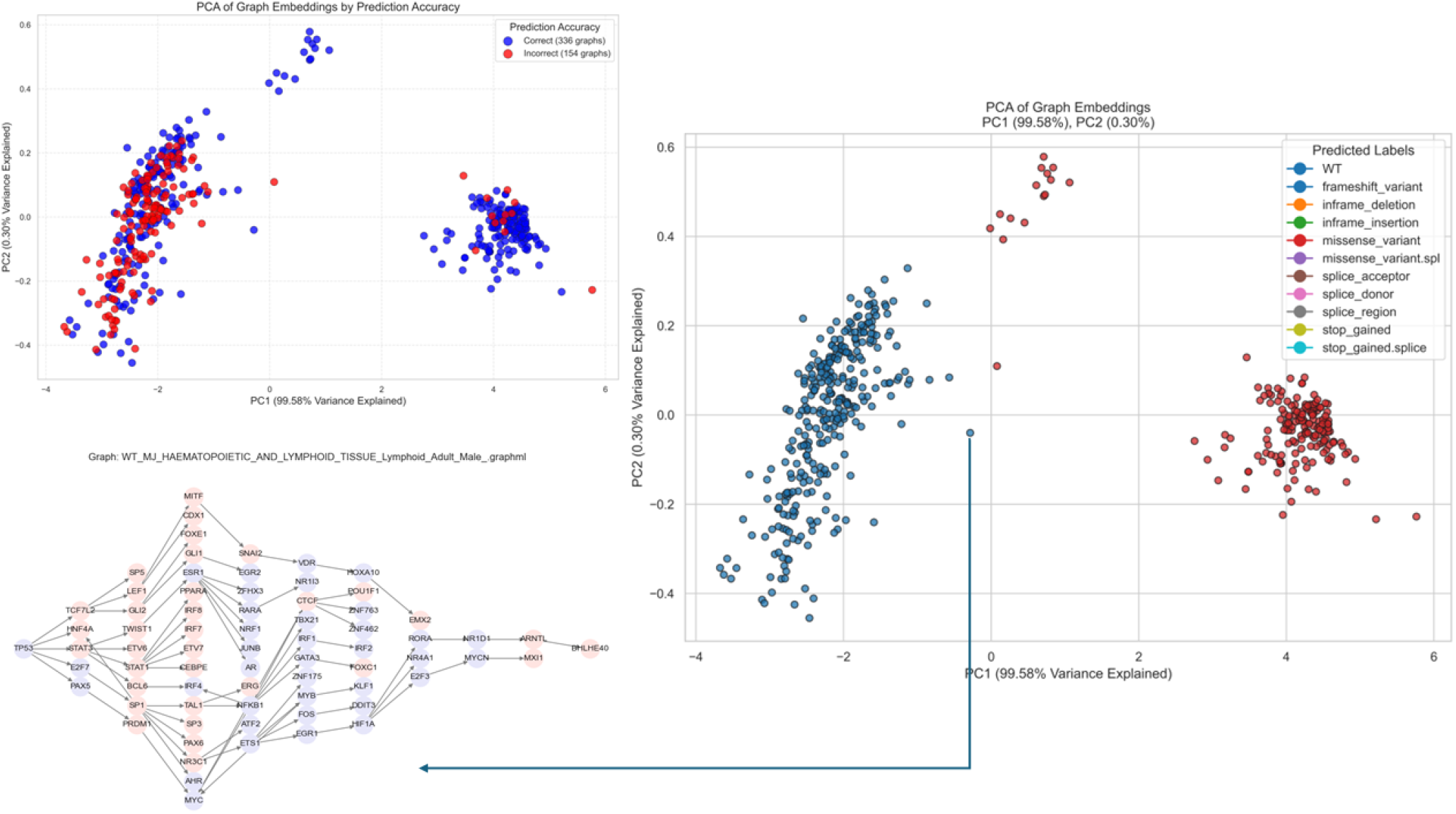
**Top left:** PCA scatter plot of the learned embeddings depicts prediction accuracy, with correctly classified graphs in blue and misclassified graphs in red, in CCLE. **Right:** PCA scatter plots of the same embeddings with different color annotation for the classified labels. Each data point maps to the actual graph/GRN (bottom left) with the gene activity profile mapped as different colors (*Lavender* activated, *Light-Rose* inactive), along with all molecular features annotated such as true mutation type, cancer site, protein change and deleterious function of mutation (if not WT). The framework maps each regulatory network as a single data-point and uses it as an input to the graph neural network classifier.

## Discussion

This study addresses the fundamental question of how to unravel the underpinnings of disease phenotypes given underlying complex gene networks and associated conditions of interest. Concretely, it shows how we can combine various methods and models for the analysis of pathway heterogeneity and ultimately resolve its association with disease states and conditions of interest. More specifically, we combined a mathematical programming approach with a deep learning model to facilitate the reconstruction and classification of gene regulatory networks that map to a specific biological state, in this case the regulon of a single *TP53* somatic mutation as a pathway of transcriptional regulation in DNA repair and damage.

Our first step was to collect a good representation of the regulon for which we used the DoRothEA collection of human regulons, including approximately 200 genes. We then extracted a starting gene regulatory PKN from *OmniPathR* to map the expression levels of the regulon and reconstruct its topology and gene activities using the mRNA sequencing from CCLE and TCGA, for each sample that carried a single *TP53* mutation. This resulted in GRNs that we then restructured via mathematical programming optimization using CARNIVAL. This framework optimizes each network per sample, given perturbations of transcription factors, and in this case we used the deleterious function of the mutation to indicate *TP53* status (active, inactive), or WT status translating into normal TF function. The optimized networks were then used as a training dataset for a Graph Neural Network which classifies them according to the type of *TP53* mutation.

The spotlight mechanism increased the classification accuracy by 16% by using *TP53* and *MYC* as our genes of interest. There may be multiple reasons behind why this synergistic set of genes works well in differentiating the regulon across different *TP53* mutations in tumor samples. These two genes share an antagonistic nature [77, 78]; *MYC* expression is significantly higher when both *MYC* amplification and *TP53* mutation are present. More specifically, *MYC* over-expression induces p53-dependent apoptosis and *TP53* mutation allows cells to tolerate supra-physiological *MYC* levels, thus promoting aggressive cancer phenotypes. This duality overrides the typical mutual exclusivity in cancer. Hence, this characteristic motif may be fitting to extract topological features, as indicated by our experiments, particularly when focusing only on the 1-hop neighborhood of these genes.

The trained classifier was able to differentiate mainly between WT and missense mutations, given the class imbalance in the dataset for the remaining MT variants, or perhaps the lack of a strong *signal* indicating heterogeneity across the other types of mutations. While the classification performance on rare mutation types was limited, this does not necessarily imply model inadequacy. Instead, it could also indicate biological similarity in downstream gene regulatory networks, where certain *TP53* mutations—even if structurally distinct—do not result in functional divergence at the transcriptional level. This is consistent with the idea that not all mutations induce measurable phenotypic effects, especially within the robust architecture of regulatory networks [76]. The classifier’ s performance, especially its difficulty in classifying rare mutations, might reflect the underlying biology, where these mutations do not produce distinct phenotypic changes. Exploratory network metrics analyzed across the two datasets (Suppl. Figures S1, S2) also did not show a strong separation of mutation types between a variety of calculated metrics. Our framework thus acts not only as a classifier but also as a lens to identify which mutations propagate sufficient regulatory alterations to be distinguishable at the network level — effectively mapping genotype-to-phenotype conversion in a data-driven manner.

While our study demonstrates the potential of integrating mathematical programming and GNNs in pathway analysis, several limitations should be taken into account. First, the size of the pathways that can be re-optimized per condition is in the order of a few hundreds of genes which raises questions about whether it is valid to incorporate these genes in a pathway without a *bona fide* knowledge of the underlying biological ground truth. Additionally, many methods exist and could be used to construct the GRNs from omics data. Furthermore, other types of GNNs and architecture settings, feature engineering, different class imbalance techniques, and oversampling strategies could be explored to potentially improve the classification performance.

We demonstrated the features and adequacy of the developed pipeline for two specific datasets looking at a specific transcription factor. However, this pipeline can be employed for any other genotype to phenotype pathway (collection of interacting genes to perform a specific biological function of interest or suspected phenotypic diversity that links to disease states), assuming the availability of data (initial PKN, expression data), and as long as the condition of interest and the disease under investigation has a strong genetic underpinning. The novelty of the approach lies in the fact that the resulting pathway variants are classified at graph-level, based on their genomic and topological characteristics: the GNN takes into account the Mode of Regulation (MoR), thus the type of interaction between the nodes (genes) as inhibition or activation, but also the gene activity profile of the pathway (the gene states of all nodes in the pathway) across other embeddings used (see Methods).

Additionally, given that networks can be a vessel of flexibility on data representation and annotation, allowing the incorporation of further genomic and transcriptomic or other (imaging) data, this framework paves the way for further expansion and inclusion of multi-modal datasets, assembling more detailed networks by the incorporation of additional features as annotated states/labels. In this case, features such as protein change, tissue origin, cancer type, gender, age, hotspot or deleterious function of mutation have been annotated on each network, as seen in Figure 6 network, as a single identifier of the network itself. This simple approach is quite flexible in expanding the array of features (clinical, molecular, etc.) used to annotate the networks, allowing straightforward change of the classification labeling scheme used in the GNN step, thus significantly increasing the possible angles of analysis. For instance, one can easily generate labels for each network based on different delimiters (e.g. protein change) and then re-train the classifier based on the new labeling scheme to now build a classifier that learns graphs that map to specific protein changes of the transcription factor (or any other gene in the network). The implementation has been tailored exactly for this type of experimentation in a modular way, allowing the GNN training and classification steps to be completely detached from the optimization step, so that it does not require repetition of GRN reconstruction, given a different labeling scheme sought. Finally, the *spotlight* mechanism (see Methods), inspired purely biologically has been effectively used to increase the attention given to specific genes of interest in the graphs. This combination of biological knowledge with state-of-the-art deep learning methods can work cooperatively and increase the efficacy of the classification. Future work could include application to different diseases or states of diseases, using different methods to generate the GRNs, classifying other conditions of interest (hotspot mutations, drug perturbations etc.) and also pruning and reconstruction of the PKNs given CRISPR [79] data (knockout experiments). These findings demonstrate the utility of graph-based representations in revealing biological signals, particularly in conditions with high molecular variability.

## Methods

*Problem Statement:* We are given a dataset 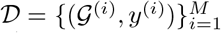 of labeled biological graphs. Each graph 𝒢= (𝒱, *ε*)represents a gene regulatory network (GRN) with:

- Nodes *v*_*i*_ *∈𝒱* corresponding to genes
- Directed edges (*i, j*) ∈ *ε* representing regulatory interactions
- Node features **x**_*i*_ ∈ ℝ^*d*^ and edge features **e**_*ij*_ ∈ ℝ^*k*^

Our goal is to learn a classifier *f* : 𝒢 ↦ *y* using a GNN, with special emphasis on a *Gene of Interest (GOI)*, such as *TP53*.

Data collected from CCLE cell-lines (n = 1,630) and TCGA tumor samples (n = 12,471) for *TP53*, incorporating either mutant/WT *TP53* samples. The DoroThEA [80] pancancer regulon was used for this transcription factor and the base PKN prior of optimization was extracted using OmniPathR [81] and accounting only for signed and directed edges. This amounted to approximately 200 downstream targets with all confidence levels included to allow the optimization step prune as appropriate and minimize the mismatch between the expression matrix and the prior knowledge network in terms of its initial topological features and mode of regulation. The CARNIVAL [82] pipeline was used to reconstruct the networks via the normalized enrichment score and the deleterious function of mutation as a perturbing event on *TP53* being the master node in all networks. The full expression matrix was used to extract the measurements for the reconstruction of the PKN for each sample, making CARNIVAL able to add or remove edges to minimize mismatch of the PKN topology and the gene activities. This mathematical programming optimization package allows for single sample reconstruction of networks and proves to be ideal in this line of work, as each network depicts exactly one CCLE model or TCGA sample carrying a single *TP53* mutation or WT state. To address class imbalance in the training data, we applied a class-weighted loss function. Specifically, class weights were computed based on the inverse frequency of each class. These weights assign a higher penalty to misclassified samples from under-represented classes. The resulting weights were then incorporated into the loss function.

### Network Reconstruction Framework (CARNIVAL)

A regulatory network is a collection of genes and their interactions; a gene can up-regulate or down-regulate other genes, and this interaction is depicted using graph theory and edges connecting nodes in a network. Each GRN was represented as an directed graph *G* = (*V, E*), where *V* represents genes (nodes) and *E* represents regulatory interactions (edges). Node attributes included gene expression and mutation status, while edge weights corresponded to the strength of regulatory interactions. Reconstructing a network amounts to the change of topology that corresponds to a particular sample (usually a full transcriptomic sequence, or else, the expression of all genes for a particular patient) so that the gene activity profiles represented by the network better fit the experimental data, than the prior knowledge. This Prior Knowledge Network (PKN) can be obtained in many different ways by combining information from multiple sources.

CARNIVAL [82] is a robust and flexible framework for causal reasoning from expression data. By integrating prior knowledge and expression-derived activities via ILP, it enables the inference of context-specific signaling networks and upstream regulators even in the absence of explicit perturbation data. Gene expression data and mutation profiles were merged and obtained from the Cancer Cell Line Encyclopedia (CCLE), alongside prior knowledge of TF-target interactions from public databases jointly processed and combined in DoRothEA. Mixed-Integer Linear Programming (MILP) is an NP-hard special case of the general Linear Programming (LP) model [83], where one imposes integrality constraints for a subset of the variables:

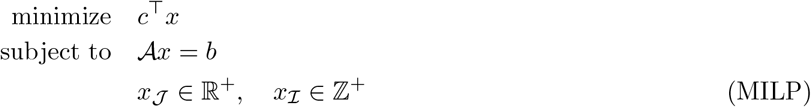

where *𝒜* ∈ ℝ^*m×n*^, *c, x* ∈ ℝ^*n*^, *b* ∈ ℝ^*m*^, *c*^⊤^ denotes the transpose of *c*, and 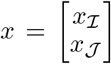. The integrality of the constraints serves as a mathematical way of mapping the prediction of link activation or removal as a binary variable. The ILP formulation seeks to minimize an objective function that balances the discrepancy between predicted and observed TF activities and the complexity of the inferred network, thus restructuring its topology, composure and directionality (MoR) to minimize loss. Constraints are imposed to ensure consistency with the prior knowledge network and the directionality and sign of interactions. For instance, if node *i* activates node *j*, and *x*_*i*_ is active, then *x*_*j*_ is constrained to be active as well. These logical relationships are translated into linear inequalities suitable for ILP solvers, along with detection of feedback loops and sparsity enforcing constraints. The MILP problem was solved using a commercial solver (IBM CPLEX) at a 5% optimality gap.

### Community Detection to extract mutation type gene signatures

Given the large number of the reconstructed networks, it proves challenging to find an efficient way of *visualizing* in a succinct way the structural differences of these networks, rendering the need for a *compression* method imperative. Our approach hence employed community detection and betweenness centrality measures to map each of these networks to a gene signature. Community detection [84] methods aim to disentangle the complexity and topology of networks, giving insights to tightly connected communities or *hubs*. Biologically, these hubs may give insight into phenotypes, identifying at the same time key nodes (genes) in the network, the gravity of which can be differently measured based on the metric used (e.g. betweenness centrality, eigenvalue centrality etc.).

We first applied an InfoMap [85] community detection algorithm, given that the graphs are directed and biological in nature, [86, 87] to split each network into tightly connected hubs/communities of nodes. We then applied betweeness centrality for each of these hubs separately, and the highest scoring genes were collected to a single signature. This technique has been used also as a node embedding feature engineering approach (see Methods). As these signatures differ even across networks carrying the same type of *TP53* mutation, we computed their intersection to form a unique for the mutation type signature. For every different mutation type a new signature hence was derived. This approach differs from [18] not only in the community detection algorithm used, but also since we now included the states of the genes (up/down) and incorporated the information in the signatures. The resulting heatmap showing each signature per mutation type, and the states of the included genes per signature can be seen in Figure 4. Collectively, these signatures convey a clear message: only missense mutations exhibit high similarity with WT samples. The rest of the variants project similarities across themselves but not with either missense, or WT samples.

### Feature Engineering

We introduced various features embedded either on nodes or edges, to fully utilize the richness of the biological regulatory network representations that CARNIVAL outputs. In this respect, nodes have activity states (up, down) and edges are not only directed but also have a MoR effect (activation, inhibition), rendering the graphs heterogeneous. These features are already encoded in the *graphml* format of the reconstructed networks from CARNIVAL and then read into graph structures which in turn dissects all node and edge features to start engineering the GNN embeddings and finally produce the associated feature matrix. A visual representation can be seen in Figure 7.

**Figure 7:**
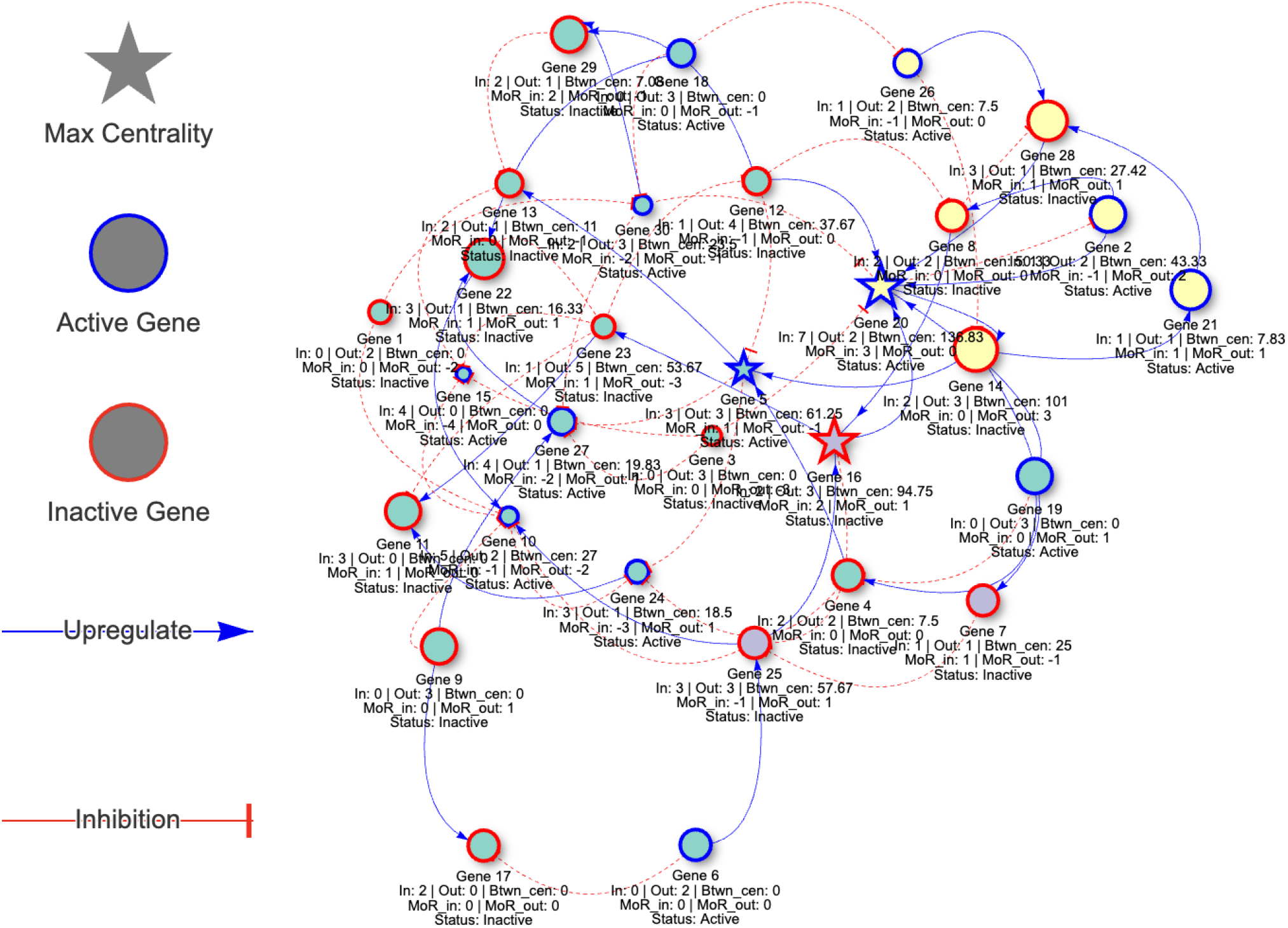
A randomly generated gene regulatory network depicted as a graph (30 nodes, 60 edges). The graph is directed and heterogeneous given that we have two different types of edges denoting inhibition and upregulation. Community detection via the Infomap algorithm dissects the graph to densely connected hubs, denoted with different node colors. For each hub we then calculate a centrality score (various metrics can be utilized here), for example the maximum betweenness centrality node here, symbolized as a star shape. Each node then has a collection of metrics calculated. The node embeddings for the graph neural network include: in degree, out degree, Status (Active/Inactive gene), centrality (within the community) and the *Split Mode of Regulation* summation which is simply the sum of all edges (1 for activating, −1 for inhibitory) across each node, as incoming and outgoing each time. The edge embeddings for the (Graph Attention Network only) are the types of the edges, again Activation (1) or Inhibition (−1), one hot encoded.

### Node Features

Each node *v*_*i*_ is associated with the following features:

- Binary color encoding: *c*_*i*_ ∈ {−1, 0, +1}
- PageRank score [88] *b*_*i*_ modulated by gene state as *b*_*i*_ * *c*_*i*_, calculated per hub from Infomap Community Detection
- In-degree 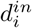 and out-degree 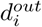
- *Split Mode of Regulation* (MoR_in_, MoR_out_)

To better capture regulatory asymmetries in the graph, we compute separate features for the cumulative incoming and outgoing regulatory effects of each node, based on the directed edges and their mode of regulation, assessing not only the regulation pressure that a node emits but also receives, captured by the *Split Mode of Regulation*. Let each edge *e* = (*u → v*) *∈ E* be associated with a regulation type: Activating (+1), Inhibitory (−1), Neutral or undefined (0). We define two node-level features, the net regulatory pressure node *v*_*i*_ either receives or exerts correspondingly:

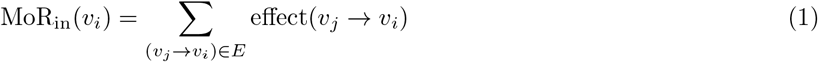

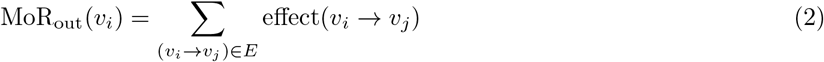

This is inspired by the potential of masking important biological distinctions. For example splitting MoR features lets the GNN differentiate such scenarios even if the node’ s degree or centrality are similar. All features (besides node color) are standardized using *z*-score normalization across the dataset. The feature vector **x** for node *v*_*i*_ becomes:

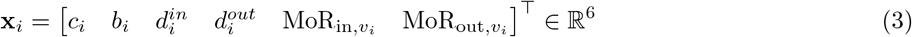

### Edge Features

Since there exist two types of edges (inhibition and activation), we sought to capture this expressiveness in the graphs to be trained by the GNN. Instead of assigning a single scalar for each edge (*i, j*) type, we assigned a one-hot vector:

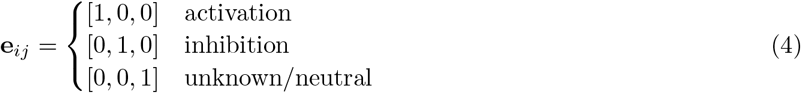

In this way, we analyzed learned edge weights to assess their contribution to the learning process (Suppl. Figure S5). A summary of the node and edge features is given in Table 1.

**Table 1:** Summary of node and edge features used in the GNN architecture.

| Feature Type | Feature | Description |
| --- | --- | --- |
| Node Features | Node Color State | Binary: 1 (lavender), -1 (mistyrose), 0 (other) |
|  | Community-weighted Centrality | PageRank score per identified community |
|  | In-degree | Number of incoming edges (scaled) |
|  | Out-degree | Number of outgoing edges (scaled) |
|  | Incoming MoR Sum | Sum of regulatory effect from incoming edges (scaled) |
|  | Outgoing MoR Sum | Sum of regulatory effect to outgoing edges (scaled) |
| Edge Features | Mode of Regulation (MoR) | One-hot: Activating [1,0,0], Inhibitory [0,1,0], Neutral [0,0,1] |

### Graph Neural Network Model

We used a Graph Attention Network (*GATv2Conv*) [20] framework consisting of attention mechanisms that learn node representations based on graph structures and edge attributes. The architecture includes multiple GATv2 convolution layers with batch normalization and residual connections. The reason for selecting this type of GNN is that it fully exploits directed and heterogeneous graphs.

Given a graph 𝒢 = (𝒱, *ε*), where 𝒱 is the set of nodes and *ε* ⊆ 𝒱 ×𝒱 is the set of edges, let:

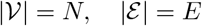

Each node *v*_*i*_ ∈ 𝒱 has an initial feature vector *x*_*i*_ ∈ ℝ^*F*^, and each edge *e*_*ij*_ ∈ *ε* has an attribute vector 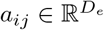. The model consists of three stacked GATv2 layers followed by a linear output layer. Each GATv2 convolution layer performs attention-based message passing with multi-head attention. For layer *l*, the embedding for node *v*_*i*_ is updated as:

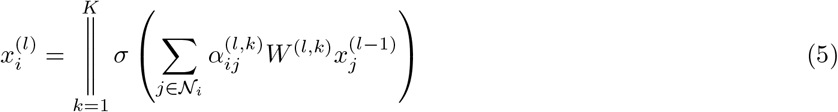

where:

- *K* is the number of attention heads.
- 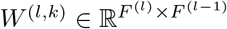is the learned weight matrix for head *k* at layer *l*.
- *σ*(*·*) is a non-linear activation (ELU in this implementation).
- ∥ denotes concatenation over all heads.
- 𝒩_*i*_ is the set of neighbors of node *v*_*i*_.

The attention coefficient 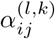 from node *v*_*j*_ to node *v*_*i*_ is calculated as:

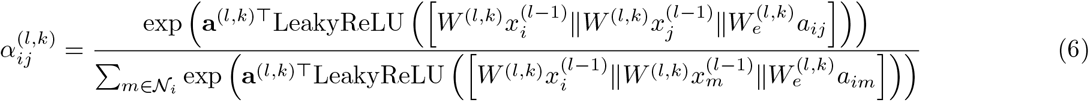

where:

- 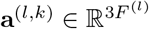 is the learned attention vector for head *k* at layer *l*. The reason 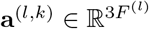 is due to the concatenation of three vectors:

- 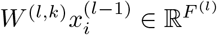, the transformed feature vector of the target node *v*_*i*_,
- 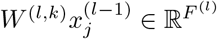, the transformed feature vector of the source node *v*_*j*_,
- 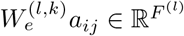, the transformed edge attributes from node *v*_*j*_ to node *v*_*i*_. Concatenating these three vectors yields a single vector in 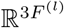, and therefore, the attention vector **a**^(*l,k*)^ must also lie in 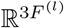 to perform the dot product operation.
- 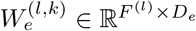 is the edge attribute transformation matrix.

The edge attributes *a*_*ij*_ are specifically modulated according to their type (activating or inhibitory). This modulation directly influences the attention coefficients, thus enhancing or suppressing message passing according to biological or functional semantics encoded in edge types. Residual connections help mitigate vanishing gradients and accelerate convergence. The output of each GATv2 layer *l* is combined with its input via a residual connection:

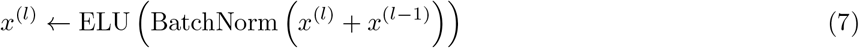

The node embeddings 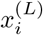 obtained from the last GATv2 layer are aggregated into a single graph-level embedding *z* through global mean pooling:

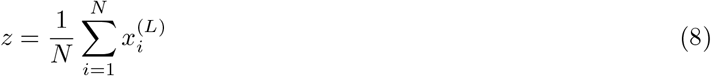

Dropout is then applied to this graph-level embedding for regularization:

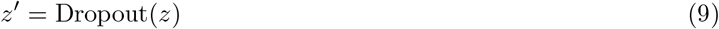

Finally, classification is performed by a linear layer:

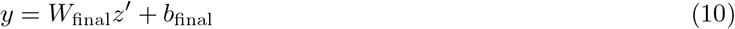

where 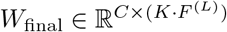 and *b*_final_ *∈* ℝ^*C*^ are the parameters of the final linear layer, with *C* being the number of classes. The loss function employed is the standard Cross-Entropy Loss for multi-class classification:

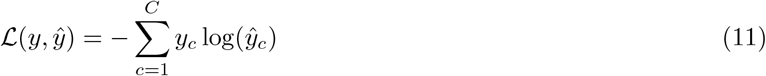

Optimization is performed using the Adam optimizer with gradient clipping to stabilize training:

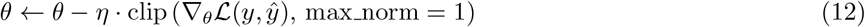

### Gene-of-Interest (GOI) Spotlight Mechanism

Since the structural nature of the input graphs represent a TF-regulon relationship (one master node regulating many target nodes), we sought to apply tailored learning that is sensitive to the close neighborhood of the TF, or a given set of genes. Given an input of a set of genes or a single gene (GOI, or *gene(s) of interest*), the implemented framework applies a spotlight mechanism (Figure 8), with scalable hop step, that can learn a mini GATv2 only for the neighbors of the GOI set, with a learnable bias vector multiplied by a learnable scalar.

**Figure 8:**
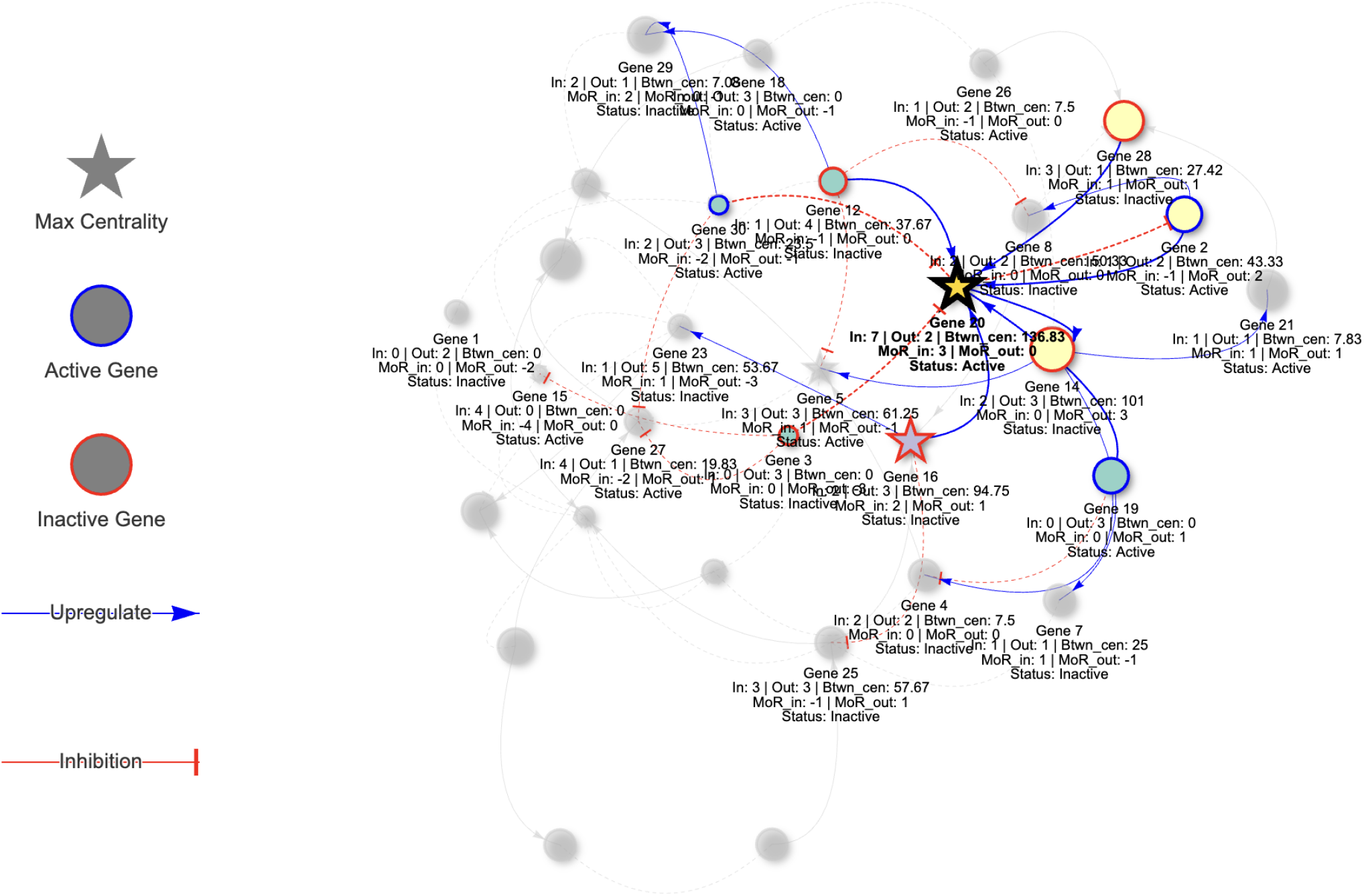
A spotlight mechanism has been implemented to allow flexibility for extra attention (learnable projection bias vector multiplied by a learnable scalar term) to a gene (or a set of genes) of interest. Given that the networks define the regulon of a transcription factor, we can assign a mini GAT layer to facilitate this biological knowledge so that the classifier is able to allow further attention to the scalable (hop modulated) neighborhood of the TF, or any other gene(s) of interest.

Let 𝒢 = (𝒱, *ε*) be a directed graph with | 𝒱 | = *N* nodes, where each node *v*_*i*_ ∈ 𝒱 is associated with a feature vector **x**_*i*_ ∈ ℝ^*d*^, and edges (*i, j*) ∈ *ε* may additionally carry attributes 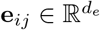. Let *𝒳 ∈* ℝ^*N* × *d*^ denote the node feature matrix, and *ε* ∈ ℝ^2× |*ε*|^ the edge index matrix. We designate a node *v*_GOI_ ∈ 𝒱 (e.g., *TP53*) as the *Gene of Interest*, or a set of nodes. We extract its *k*-hop neighborhood:

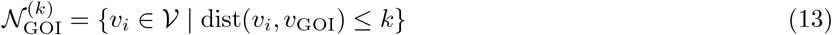

and the induced subgraph 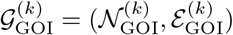, where 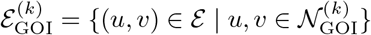. We apply a dedicated multi-head Graph Attention Network (GAT) layer to this subgraph:

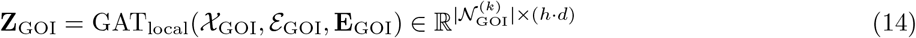

where *h* is the number of attention heads. To maintain dimensional consistency with the global graph, we project the local embeddings:

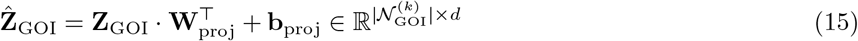

Additionally, we introduce a learnable bias vector **b**_GOI_ *∈* ℝ^*d*^ and a scalar *α*_GOI_ *∈*ℝ to enhance the GOI neighborhood:

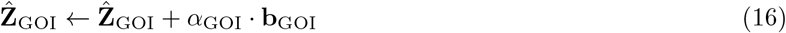

Finally, we overwrite the updated GOI neighborhood features into the global node feature matrix:

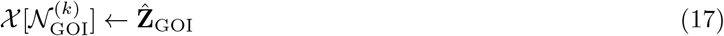

The updated node features are processed through a standard stack of *L* GATv2 layers with residual connections:

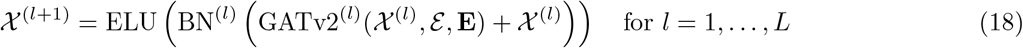

The final graph embedding is obtained by mean pooling and passed through a linear classifier:

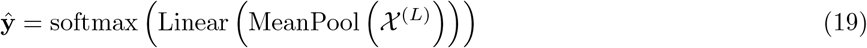

### Hyper-Parameter Tuning

We performed automated hyper-parameter optimization of our GNN using the Optuna framework [89]. The search targeted minimization of the misclassification rate on a test set, using a Tree-structured Parzen Estimator (TPE) as the sampling algorithm. The objective function called a GNN training pipeline with a set of hyper-parameters sampled from predefined spaces. The train-test split ratio was set to 0.7 for both datasets with label balancing.

At the conclusion of the study, we extracted the best hyper-parameter set and generated a static plot showing model performance across 50 trials (see Suppl. Figure S6). The computational tests were performed on an M4 MacBookAir Processor with 24GB of RAM, rendering our approach feasible *at-laptop-scale*. Optimal configurations can be found in Table 2.

**Table 2:** Hyperparameter search space and best values selected by Optuna for TCGA and CCLE datasets.

| Parameter | Distribution | Search Space | TCGA-Best | CCLE-Best |
| --- | --- | --- | --- | --- |
| Hidden Dimension | Categorical | {32, 64} | 64 | 32 |
| Dropout Probability | Uniform Float | [0.2, 0.5] | 0.23 | 0.38 |
| Learning Rate | Log-Uniform Float | $[10^{-5}, 10^{-3}]$ | $2.47 \times 10^{-4}$ | $6.21 \times 10^{-5}$ |
| Number of Attention Heads | Discrete Integer | {2, 4, 6} | 2 | 2 |
| Number of Epochs | Categorical | {50, 100} | 100 | 50 |
| Batch Size | Categorical | {16, 32, 64} | 16 | 16 |
| GOI Number of Hops | Discrete Integer | {1, 2, 3} | 1 | 1 |

## Acknowledgments

We wish to thank the anonymous reviewers for their valuable comments and feedback that helped improve the quality of the manuscript.

## Author Contributions

Conceptualization: C.P.T.

Methodology: C.P.T., R.A.

Software: C.P.T.

Formal analysis: C.P.T.

Investigation: C.P.T., R.A.

Writing – original draft: C.P.T.

Writing – review & editing: R.A.

Visualization: C.P.T.

Supervision: C.P.T., R.A.

## Competing interests

The authors declare no competing interests.

## Data Availability

The repository associated with this work can be found in https://github.com/harrytr/HarmonizeR, where we built a Shiny R application with user interface to facilitate reproducibility of our experiments. For cell lines, data have been collected from DepMap 24Q2 https://depmap.org/portal/data_page/?tab=currentRelease. For tumor samples, the TCGA dataset has been collected along with known mutation profiles from https://gdc.cancer.gov/about-data/publications/pancanatlas. Data from DoroThEA was collected using OmniPathR (using import of DoroThEA interactions command for human organism) and extracting only directed and signed edges.

## Notes

https://github.com/harrytr/HarmonizeR

### Competing Interest Statement

The authors have declared no competing interest.

### Summary of Updates

-We changed the title -We now demonstrate its generalizability by also applying it to the TCGA (tumour samples) dataset. -The ANM step was removed. -From using the Convolutional GNN, we moved to using the GATv2Conv model -We implemented a spotlight mechanism -We added automated hyperparameter tuning for our GNN

## References

[1] Dominick A Rizzi and Stig Andur Pedersen. Causality in medicine: towards a theory and terminology. Theoretical Medicine, 13:233–254, 1992.

[2] Jean-François Bach. Causality in medicine. Comptes Rendus. Biologies, 342(3-4):55–57, 2019.

[3] Jonathan G Richens, Ciarán M Lee, and Saurabh Johri. Improving the accuracy of medical diagnosis with causal machine learning. Nature communications, 11(1):3923, 2020.

[4] Gajendra Jung Katuwal and Robert Chen. Machine learning model interpretability for precision medicine. arXiv preprint 1610.09045, 2016.

[5] Leilani H Gilpin, David Bau, Ben Z Yuan, Ayesha Bajwa, Michael Specter, and Lalana Kagal. Explaining explanations: An overview of interpretability of machine learning. In 2018 IEEE 5th International Conference on data science and advanced analytics (DSAA), pages 80–89. IEEE, 2018.

[6] Diogo V Carvalho, Eduardo M Pereira, and Jaime S Cardoso. Machine learning interpretability: A survey on methods and metrics. Electronics, 8(8):832, 2019.

[7] Alfredo Vellido. The importance of interpretability and visualization in machine learning for applications in medicine and health care. Neural computing and applications, 32(24):18069–18083, 2020.

[8] Paul Smolen, Douglas A Baxter, and John H Byrne. Mathematical modeling of gene networks. Neuron, 26(3):567–580, June 2000.

[9] Paul Brazhnik, Alberto de la Fuente, and Pedro Mendes. Gene networks: how to put the function in genomics. Trends in Biotechnology, 20(11):467–472, November 2002.

[10] Richard Bonneau. Learning biological networks: from modules to dynamics. Nature Chemical Biology, 4(11):658–664, November 2008.

[11] Guy Karlebach and Ron Shamir. Modelling and analysis of gene regulatory networks. Nature Reviews Molecular Cell Biology, 9(10):770–780, October 2008.

[12] Ernst Althaus, Gunnar W. Klau, Oliver Kohlbacher, Hans-Peter Lenhof, and Knut Reinert. Integer Linear Programming in Computational Biology, pages 199–218. Springer Berlin Heidelberg, Berlin, Heidelberg, 2009.

[13] Nicolas Le Novére. Quantitative and logic modelling of molecular and gene networks. Nature Reviews Genetics, 16(3):146–158, March 2015.

[14] J. Pearl, M. Glymour, and N.P. Jewell. Causal Inference in Statistics: A Primer. Wiley, 2016.

[15] Guy Karlebach and Ron Shamir. Modelling and analysis of gene regulatory networks. Nature reviews Molecular cell biology, 9(10):770–780, 2008.

[16] Douglas H Erwin and Eric H Davidson. The evolution of hierarchical gene regulatory networks. Nature Reviews Genetics, 10(2):141–148, 2009.

[17] D. Bertsimas and J.N. Tsitsiklis. Introduction to linear optimization. Athena Scientific, 1997.

[18] Charalampos P. Triantafyllidis, Alessandro Barberis, Fiona Hartley, Ana Miar Cuervo, Enio Gjerga, Philip Charlton, Linda van Bijsterveldt, Julio Saez Rodriguez, and Francesca M. Buffa. A machine learning and directed network optimization approach to uncover tp53 regulatory patterns. iScience, 26(12), Dec 2023.

[19] Aviad Tsherniak, Francisca Vazquez, Phil G Montgomery, Barbara A Weir, Gregory Kryukov, Glenn S Cowley, Stanley Gill, William F Harrington, Sasha Pantel, John M Krill-Burger, et al. Defining a cancer dependency map. Cell, 170(3):564–576, 2017.

[20] Shaked Brody, Uri Alon, and Eran Yahav. How attentive are graph attention networks?, 2022.

[21] Soo Min Lee, Younghyun Han, and Kwang-Hyun Cho. Deep learning untangles the resistance mechanism of p53 reactivator in lung cancer cells. iScience, 26(12), Dec 2023.

[22] Hong Ling, Sandhya Samarasinghe, and Don Kulasiri. Novel recurrent neural network for modelling biological networks: Oscillatory p53 interaction dynamics. Biosystems, 114(3):191–205, 2013.

[23] Shuyu Wang, Hongzhou Tang, Peng Shan, Zhaoxia Wu, and Lei Zuo. Pros-gnn: Predicting effects of mutations on protein stability using graph neural networks. Computational Biology and Chemistry, 107:107952, 2023.

[24] Hao Li, Zebei Han, Yu Sun, Fu Wang, Pengzhen Hu, Yuang Gao, Xuemei Bai, Shiyu Peng, Chao Ren, Xiang Xu, Zeyu Liu, Hebing Chen, Yang Yang, and Xiaochen Bo. Cgmega: explainable graph neural network framework with attention mechanisms for cancer gene module dissection. Nature Communications, 15(1):5997, Jul 2024.

[25] Gil Ben Cohen, Adar Yaacov, Yishai Ben Zvi, Ranel Loutati, Natan Lishinsky, Jakob Landau, Tom Hope, Aron Popovzter, and Shai Rosenberg. Graph convolution networks model identifies and quantifies gene and cancer specific transcriptome signatures of cancer driver events. Computers in Biology and Medicine, 185:109491, 2025.

[26] Narumi Hatano, Mayumi Kamada, Ryosuke Kojima, and Yasushi Okuno. Network-based prediction approach for cancer-specific driver missense mutations using a graph neural network. BMC Bioinformatics, 24(1):383, October 2023.

[27] Thomas N Kipf and Max Welling. Semi-supervised classification with graph convolutional networks. arXiv preprint 1609.02907, 2016.

[28] Gabriele Corso, Hannes Stark, Stefanie Jegelka, Tommi Jaakkola, and Regina Barzilay. Graph neural networks. Nature Reviews Methods Primers, 4(1):17, 2024.

[29] William L Hamilton. Graph representation learning. Morgan & Claypool Publishers, 2020.

[30] Zonghan Wu, Shirui Pan, Fengwen Chen, Guodong Long, Chengqi Zhang, and S Yu Philip. A comprehensive survey on graph neural networks. IEEE transactions on neural networks and learning systems, 32(1):4–24, 2020.

[31] Xiao-Meng Zhang, Li Liang, Lin Liu, and Ming-Jing Tang. Graph neural networks and their current applications in bioinformatics. Frontiers in Genetics, Volume 12-2021, 2021.

[32] Sijie Li, Heyang Hua, and Shengquan Chen. Graph neural networks for single-cell omics data: a review of approaches and applications. Brief Bioinform, 26(2), March 2025.

[33] Emma Paul M., Jereesh A.S., and G. Santhosh Kumar. Reconstruction of gene regulatory networks using graph neural networks. Applied Soft Computing, 163:111899, 2024.

[34] Michail Chatzianastasis, Michalis Vazirgiannis, and Zijun Zhang. Explainable multilayer graph neural network for cancer gene prediction. Bioinformatics, 39(11):btad643, 10 2023.

[35] Lingling Zhang, Shaowei Wang, Jun Liu, Xiaojun Chang, Qika Lin, Yaqiang Wu, and Qinghua Zheng. Mul-grn: Multi-level graph relation network for few-shot node classification. IEEE Transactions on Knowledge and Data Engineering, 35(6):6085–6098, 2023.

[36] Guo Mao, Zhengbin Pang, Ke Zuo, Qinglin Wang, Xiangdong Pei, Xinhai Chen, and Jie Liu. Predicting gene regulatory links from single-cell rna-seq data using graph neural networks. Briefings in Bioinformatics, 24(6):bbad414, 11 2023.

[37] Xiaohan Xing, Fan Yang, Hang Li, Jun Zhang, Yu Zhao, Mingxuan Gao, Junzhou Huang, and Jianhua Yao. Multi-level attention graph neural network based on co-expression gene modules for disease diagnosis and prognosis. Bioinformatics, 38(8):2178–2186, 02 2022.

[38] Rui Li, Xin Yuan, Mohsen Radfar, Peter Marendy, Wei Ni, Terrence J. O’Brien, and Pablo M. Casillas-Espinosa. Graph signal processing, graph neural network and graph learning on biological data: A systematic review. IEEE Reviews in Biomedical Engineering, 16:109–135, 2023.

[39] Yong Li, Xiao Song, Kaiqi Gong, Songsong Liu, and Wenxin Li. Differentially private graph neural networks for graph classification and its adaptive optimization. Expert Systems with Applications, 263:125798, 2025.

[40] Pengbo Duan, Kuo Yang, Xin Su, Shuyue Fan, Xin Dong, Fenghui Zhang, Xianan Li, Xiaoyan Xing, Qiang Zhu, Jian Yu, and Xuezhong Zhou. Htinet2: herb–target prediction via knowledge graph embedding and residual-like graph neural network. Briefings in Bioinformatics, 25(5):bbae414, 08 2024.

[41] Juexin Wang, Anjun Ma, Yuzhou Chang, Jianting Gong, Yuexu Jiang, Ren Qi, Cankun Wang, Hongjun Fu, Qin Ma, and Dong Xu. scgnn is a novel graph neural network framework for single-cell rna-seq analyses. Nature Communications, 12(1):1882, Mar 2021.

[42] Tianyu Liu, Yuge Wang, Zhitao Ying, and Hongyu Zhao. Muse-GNN: Learning unified gene representation from multimodal biological graph data. In Thirty-seventh Conference on Neural Information Processing Systems, 2023.

[43] Rajat Mishra and S. Shridevi. Knowledge graph driven medicine recommendation system using graph neural networks on longitudinal medical records. Scientific Reports, 14(1):25449, Oct 2024.

[44] Chengsheng Mao, Liang Yao, and Yuan Luo. Medgcn: Medication recommendation and lab test imputation via graph convolutional networks. Journal of Biomedical Informatics, 127:104000, 2022.

[45] Ye Yuan and Ziv Bar-Joseph. Gcng: graph convolutional networks for inferring gene interaction from spatial transcriptomics data. Genome Biology, 21(1):300, Dec 2020.

[46] Bilin Liang, Haifan Gong, Lu Lu, and Jie Xu. Risk stratification and pathway analysis based on graph neural network and interpretable algorithm. BMC Bioinformatics, 23(1):394, Sep 2022.

[47] András Gézsi and Péter Antal. Gnn4dm: a graph neural network-based method to identify overlapping functional disease modules. Bioinformatics, 40(10):btae573, 09 2024.

[48] Pi-Jing Wei, Ziqiang Guo, Zhen Gao, Zheng Ding, Rui-Fen Cao, Yansen Su, and Chun-Hou Zheng. Inference of gene regulatory networks based on directed graph convolutional networks. Briefings in Bioinformatics, 25(4):bbae309, 06 2024.

[49] Hongxi Yan, Dawei Weng, Dongguo Li, Yu Gu, Wenji Ma, and Qingjie Liu. Prior knowledge-guided multilevel graph neural network for tumor risk prediction and interpretation via multi-omics data integration. Briefings in Bioinformatics, 25(3):bbae184, 04 2024.

[50] Douglas Hanahan. Hallmarks of cancer: New dimensions. Cancer Discov, 12(1):31–46, January 2022.

[51] Amaia Lujambio and Scott W Lowe. The microcosmos of cancer. Nature, 482(7385):347–355, 2012.

[52] Douglas Hanahan and Robert A Weinberg. The hallmarks of cancer. cell, 100(1):57–70, 2000.

[53] Douglas Hanahan and Robert A Weinberg. Hallmarks of cancer: the next generation. cell, 144(5):646–674, 2011.

[54] Chadd E Nesbit, Jean M Tersak, and Edward V Prochownik. Myc oncogenes and human neoplastic disease. Oncogene, 18(19):3004–3016, 1999.

[55] Samuel A Lambert, Arttu Jolma, Laura F Campitelli, Pratyush K Das, Yimeng Yin, Mihai Albu, Xiaoting Chen, Jussi Taipale, Timothy R Hughes, and Matthew T Weirauch. The human transcription factors. Cell, 172(4):650–665, 2018.

[56] Dimas Yusuf, Stefanie L Butland, Magdalena I Swanson, Eugene Bolotin, Amy Ticoll, Warren A Cheung, Xiao Yu Cindy Zhang, Christopher TD Dickman, Debra L Fulton, Jonathan S Lim, et al. The transcription factor encyclopedia. Genome biology, 13:1–25, 2012.

[57] Athanasios G Papavassiliou. Transcription factors. New England Journal of Medicine, 332(1):45–47, 1995.

[58] Henry E Pratt, Gregory R Andrews, Nishigandha Phalke, Michael J Purcaro, Arjan van der Velde, Jill E Moore, and Zhiping Weng. Factorbook: an updated catalog of transcription factor motifs and candidate regulatory motif sites. Nucleic Acids Res, 50(D1):D141–D149, January 2022.

[59] William R Wilson and Michael P Hay. Targeting hypoxia in cancer therapy. Nature Reviews Cancer, 11(6):393–410, 2011.

[60] J Martin Brown and William R Wilson. Exploiting tumour hypoxia in cancer treatment. Nature Reviews Cancer, 4(6):437–447, 2004.

[61] T Soussi and KG Wiman. Tp53: an oncogene in disguise. Cell Death & Differentiation, 22(8):1239–1249, 2015.

[62] Magali Olivier, Monica Hollstein, and Pierre Hainaut. Tp53 mutations in human cancers: origins, consequences, and clinical use. Cold Spring Harbor perspectives in biology, 2(1):a001008, 2010.

[63] Achatz Petitjean, MIW Achatz, AL Borresen-Dale, P Hainaut, and M Olivier. Tp53 mutations in human cancers: functional selection and impact on cancer prognosis and outcomes. Oncogene, 26(15):2157–2165, 2007.

[64] Monica Hollstein, David Sidransky, Bert Vogelstein, and Curtis C Harris. p53 mutations in human cancers. Science, 253(5015):49–53, 1991.

[65] Pierre May and Evelyne May. Twenty years of p53 research: structural and functional aspects of the p53 protein. Oncogene, 18(53):7621–7636, 1999.

[66] David P Lane. p53, guardian of the genome. Nature, 358(6381), 1992.

[67] Cristian Tomasetti, Lu Li, and Bert Vogelstein. Stem cell divisions, somatic mutations, cancer etiology, and cancer prevention. Science, 355(6331):1330–1334, 2017.

[68] Lawrence A. Donehower, Thierry Soussi, Anil Korkut, Yuexin Liu, Andre Schultz, Maria Cardenas, Xubin Li, Ozgun Babur, Teng-Kuei Hsu, Olivier Lichtarge, John N. Weinstein, Rehan Akbani, and David A. Wheeler. Integrated analysis of tp53 gene and pathway alterations in the cancer genome atlas. Cell Reports, 28(5):1370 – 1384.e5, 2019.

[69] Jan-Philipp Kruse and Wei Gu. Modes of p53 regulation. Cell, 137(4):609–622, 2009.

[70] Xiaohua Chen, Taotao Zhang, Wei Su, Zhihui Dou, Dapeng Zhao, Xiaodong Jin, Huiwen Lei, Jing Wang, Xiaodong Xie, Bo Cheng, Qiang Li, Hong Zhang, and Cuixia Di. Mutant p53 in cancer: from molecular mechanism to therapeutic modulation. Cell Death & Disease, 13(11):974, Nov 2022.

[71] Andreas C Joerger, Thorsten Stiewe, and Thierry Soussi. TP53: the unluckiest of genes? Cell Death Differ, 32(2):219–224, October 2024.

[72] Andreas C Joerger, Thorsten Stiewe, and Thierry Soussi. Tp53: the unluckiest of genes? Cell Death & Differentiation, pages 1–6, 2024.

[73] Gizem Efe, Anil K Rustgi, and Carol Prives. p53 at the crossroads of tumor immunity. Nature Cancer, 5(7):983–995, July 2024.

[74] Qisheng Pan, Stephanie Portelli, Thanh Binh Nguyen, and David B Ascher. Characterization on the oncogenic effect of the missense mutations of p53 via machine learning. Briefings in Bioinformatics, 25(1):bbad428, 11 2023.

[75] Gil Ben-Cohen, Flora Doffe, Michal Devir, Bernard Leroy, Thierry Soussi, and Shai Rosenberg. Tp53 prof: a machine learning model to predict impact of missense mutations in tp53. Briefings in Bioinformatics, 23(2):bbab524, 01 2022.

[76] Julianne S. Funk, Maria Klimovich, Daniel Drangenstein, Ole Pielhoop, Pascal Hunold, Anna Borowek, Maxim Noeparast, Evangelos Pavlakis, Michelle Neumann, Dimitrios-Ilias Balourdas, Katharina Kochhan, Nastasja Merle, Imke Bullwinkel, Michael Wanzel, Sabrina Elmshaüser, Julia Teply-Szymanski, Andrea Nist, Tara Procida, Marek Bartkuhn, Katharina Humpert, Marco Mernberger, Rajkumar Savai, Thierry Soussi, Andreas C. Joerger, and Thorsten Stiewe. Deep crispr mutagenesis characterizes the functional diversity of tp53 mutations. Nature Genetics, 57(1):140–153, Jan 2025.

[77] Peter Ulz, Ellen Heitzer, and Michael R. Speicher. Co-occurrence of myc amplification and tp53 mutations in human cancer. Nature Genetics, 48(2):104–106, Feb 2016.

[78] Sander Canisius, John W. M. Martens, and Lodewyk F. A. Wessels. A novel independence test for somatic alterations in cancer shows that biology drives mutual exclusivity but chance explains most co-occurrence. Genome Biology, 17(1):261, Dec 2016.

[79] Prashant Mali, Luhan Yang, Kevin M Esvelt, John Aach, Marc Guell, James E DiCarlo, Julie E Norville, and George M Church. RNA-guided human genome engineering via cas9. Science, 339(6121):823–826, January 2013.

[80] Luz Garcia-Alonso, Christian H Holland, Mahmoud M Ibrahim, Denes Turei, and Julio Saez-Rodriguez. Benchmark and integration of resources for the estimation of human transcription factor activities. Genome research, 29(8):1363–1375, 2019.

[81] Denes Turei, Tamas Korcsmaros, and Julio Saez-Rodriguez. Omnipath: Guidelines and gateway for literature-curated signaling pathway resources. Nature Methods, 13:966–967, 11 2016.

[82] Anika Liu, Panuwat Trairatphisan, Enio Gjerga, Athanasios Didangelos, Jonathan Barratt, and Julio Saez-Rodriguez. From expression footprints to causal pathways: contextualizing large signaling networks with carnival. npj Systems Biology and Applications, 5(1):40, 2019.

[83] Charalampos P Triantafyllidis and Nikolaos Samaras. A new non-monotonic infeasible simplex-type algorithm for linear programming. PeerJ Comput Sci, 6:e265, March 2020.

[84] M. E. J. Newman and M. Girvan. Finding and evaluating community structure in networks. Phys. Rev. E, 69:026113, Feb 2004.

[85] Martin Rosvall and Carl T. Bergstrom. Maps of random walks on complex networks reveal community structure. Proceedings of the National Academy of Sciences, 105(4):1118–1123, 2008.

[86] Jelena Smiljanić, Christopher Blöcker, Anton Holmgren, Daniel Edler, Magnus Neuman, and Martin Rosvall. Community detection with the map equation and infomap: Theory and applications, 2023.

[87] Martin Rosvall and Carl T. Bergstrom. Maps of random walks on complex networks reveal community structure. Proceedings of the National Academy of Sciences, 105(4):1118–1123, 2008.

[88] Lawrence Page, Sergey Brin, Rajeev Motwani, and Terry Winograd. The pagerank citation ranking: Bringing order to the web. Technical Report 1999-66, Stanford InfoLab, November 1999. Previous number = SIDL-WP-1999-0120.

[89] Takuya Akiba, Shotaro Sano, Toshihiko Yanase, Takeru Ohta, and Masanori Koyama. Optuna: A next-generation hyperparameter optimization framework. In Proceedings of the 25th ACM SIGKDD International Conference on Knowledge Discovery & Data Mining, KDD ‘19, page 2623–2631, New York, NY, USA, 2019. Association for Computing Machinery.

