## Supplementary material for "Causality-aware graph neural networks for functional stratification and phenotype prediction at scale": SI

Supplemental Information

Charalampos P. Triantafyllidis*
Nuffield Department of Medicine,
University of Oxford, Oxford, OX3 7BN, UK


Ricardo Aguas
Nuffield Department of Medicine,
University of Oxford, Oxford, OX3 7BN, UK


CCLE GNN Classifier evaluation:

| | precision | recall | f1-score | support |
|---|---|---|---|---|
| WT | 0.55 | 0.96 | 0.70 | 179 |
| frameshift_variant | 0.00 | 0.00 | 0.00 | 33 |
| inframe_deletion | 0.00 | 0.00 | 0.00 | 6 |
| inframe_insertion | 0.00 | 0.00 | 0.00 | 1 |
| missense_variant | 0.93 | 0.82 | 0.87 | 201 |
| missense_variant.spl | 0.00 | 0.00 | 0.00 | 5 |
| splice_acceptor | 0.00 | 0.00 | 0.00 | 14 |
| splice_donor | 0.00 | 0.00 | 0.00 | 12 |
| splice_region | 0.00 | 0.00 | 0.00 | 4 |
| stop_gained | 0.00 | 0.00 | 0.00 | 33 |
| stop_gained.splice | 0.00 | 0.00 | 0.00 | 2 |
| accuracy | | | 0.69 | 490 |
| macro avg | 0.13 | 0.16 | 0.14 | 490 |
| weighted avg | 0.58 | 0.69 | 0.61 | 490 |

TCGA GNN Classifier evaluation:

| | precision | recall | f1-score | support |
|---|---|---|---|---|
| Frame_Shift_Del | 0.00 | 0.00 | 0.00 | 117 |
| Frame_Shift_Ins | 0.00 | 0.00 | 0.00 | 36 |
| Fusion | 0.00 | 0.00 | 0.00 | 8 |
| In_Frame_Del | 0.00 | 0.00 | 0.00 | 23 |
| In_Frame_Ins | 0.00 | 0.00 | 0.00 | 4 |
| Missense_Mutation | 0.99 | 0.83 | 0.90 | 808 |
| Nonsense_Mutation | 0.00 | 0.00 | 0.00 | 164 |
| Splice_Region | 0.00 | 0.00 | 0.00 | 15 |
| Splice_Site | 0.00 | 0.00 | 0.00 | 83 |
| WT | 0.81 | 1.00 | 0.89 | 2484 |
| accuracy | | | 0.84 | 3742 |
| macro avg | 0.18 | 0.18 | 0.18 | 3742 |
| weighted avg | 0.75 | 0.84 | 0.79 | 3742 |

TCGA GNN Classifier evaluation (without GOI spotlight):

```
                   precision    recall  f1-score   support

  Frame_Shift_Del       0.05      0.01      0.01       117
  Frame_Shift_Ins       0.00      0.00      0.00        36
           Fusion       0.00      0.00      0.00         8
     In_Frame_Del       0.00      0.00      0.00        23
     In_Frame_Ins       0.00      0.00      0.00         4
Missense_Mutation       1.00      0.83      0.91       808
Nonsense_Mutation       0.08      0.46      0.14       164
    Splice_Region       0.00      0.00      0.00        15
      Splice_Site       0.00      0.00      0.00        83
               WT       0.85      0.72      0.78      2484

         accuracy                           0.68      3742
        macro avg       0.20      0.20      0.18      3742
     weighted avg       0.78      0.68      0.72      3742

Confusion Matrix:
 [[1781    0    0   13    2    0  688    0    0    0]
 [  17    0    0    2    0    0   17    0    0    0]
 [  15    0    0    1    0    0    7    0    0    0]
 [  66    0    0    1    0    0   50    0    0    0]
 [  75    0    0    0  672    0   61    0    0    0]
 [   8    0    0    0    0    0    0    0    0    0]
 [  85    0    0    3    0    0   76    0    0    0]
 [   2    0    0    0    0    0    2    0    0    0]
 [   7    0    0    0    0    0    8    0    0    0]
 [  49    0    0    0    0    0   34    0    0    0]]
Total Graphs: 3742
Misclassified Graphs: 1212
Misclassification Percentage: 32.39%
```

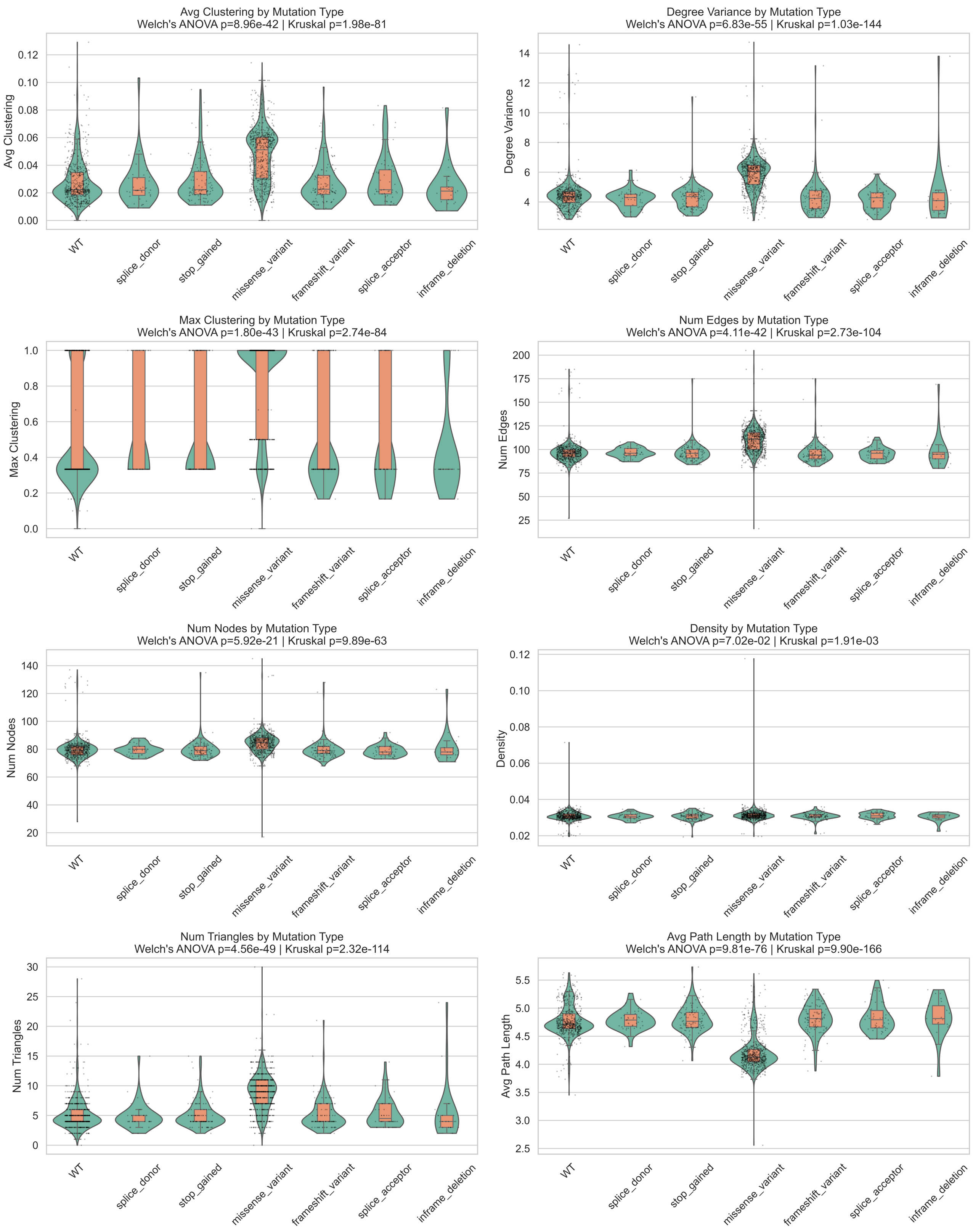


Figure S1: CCLE

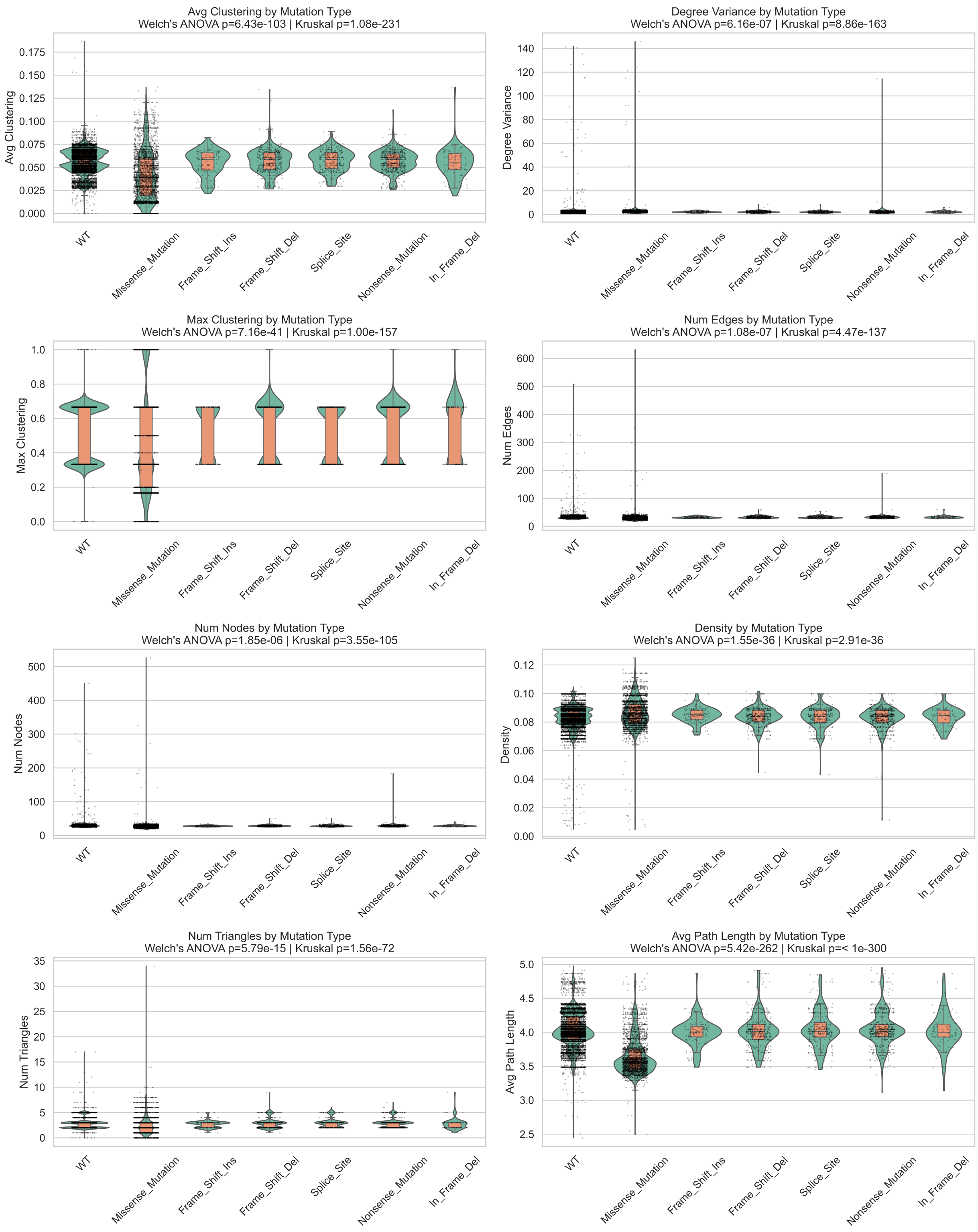


Figure S2: TCGA

Figures S1 and S2: Analyzing various network metrics (average clustering, degree variance, max clustering, number of edges and nodes, density, number of triangles, and average path length) across the most prevalent mutation types. Significant differences are observed across WT and missense samples mainly.

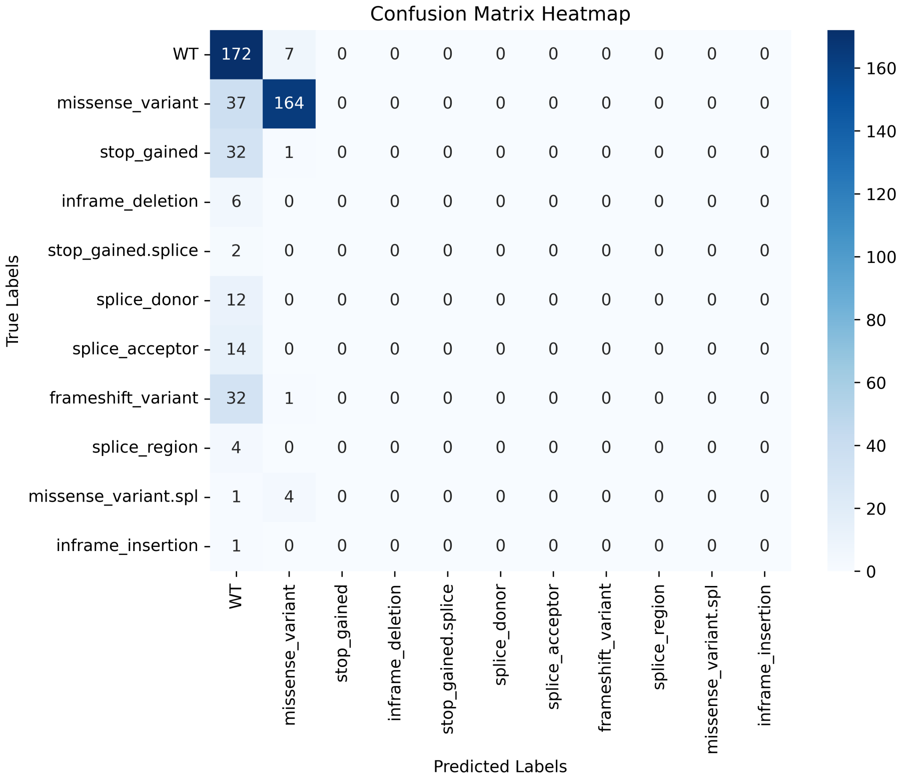


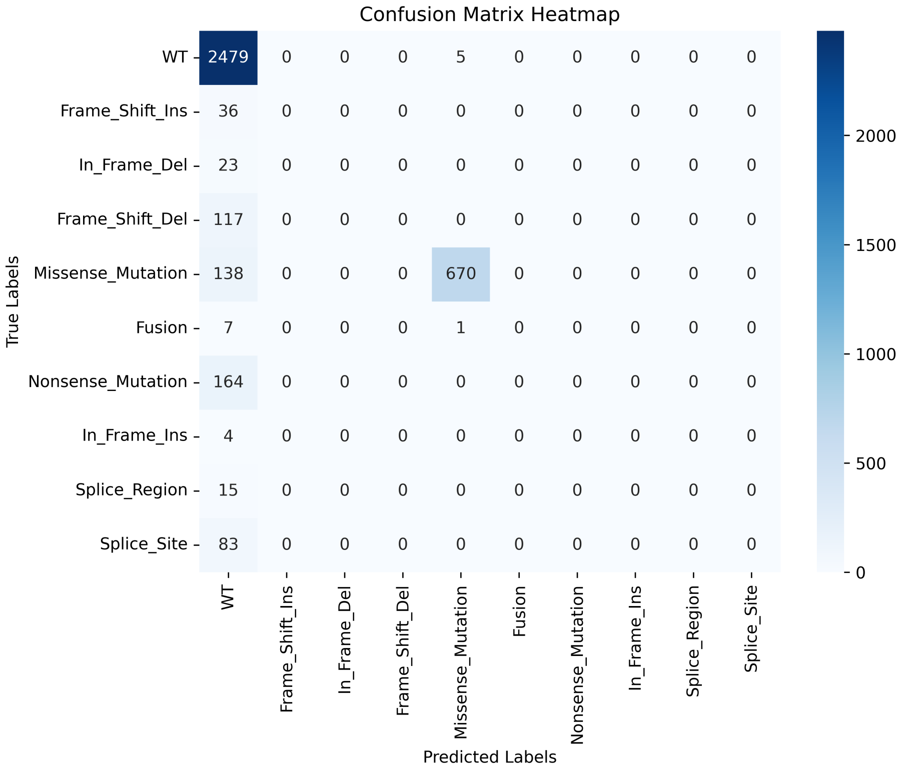


Figure S3: Confusion Matrix for the multi-label classifier built on GNNs (CCLE top, TCGA bottom). The classifier, as expected given the class imbalance and the availability of different types of mutations for *TP53* detects very well missense and WT samples (the most prevalent ones), and poorly other types.

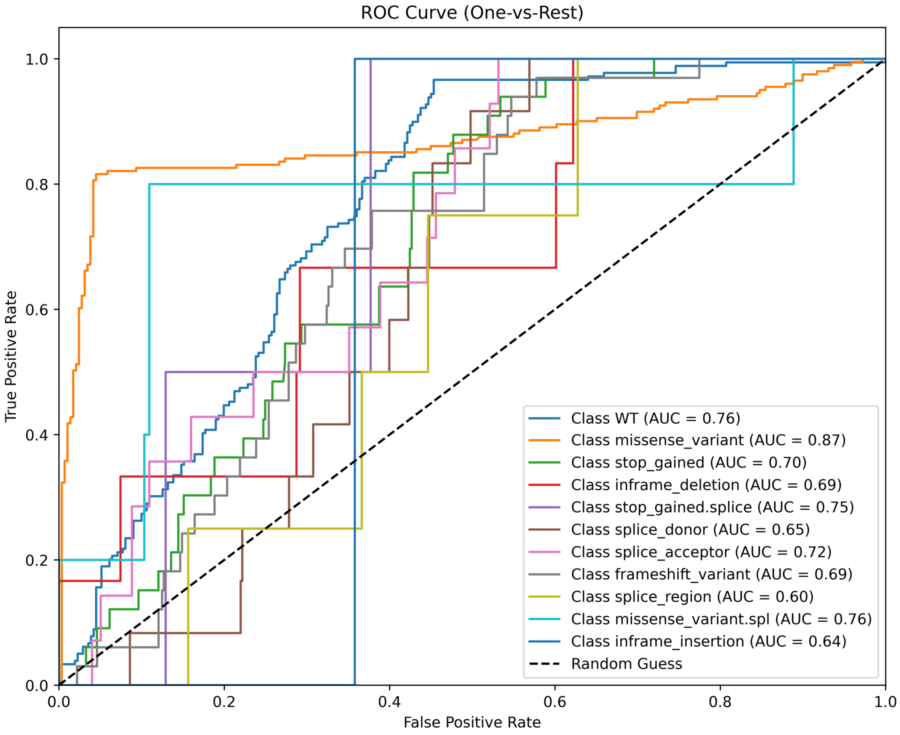


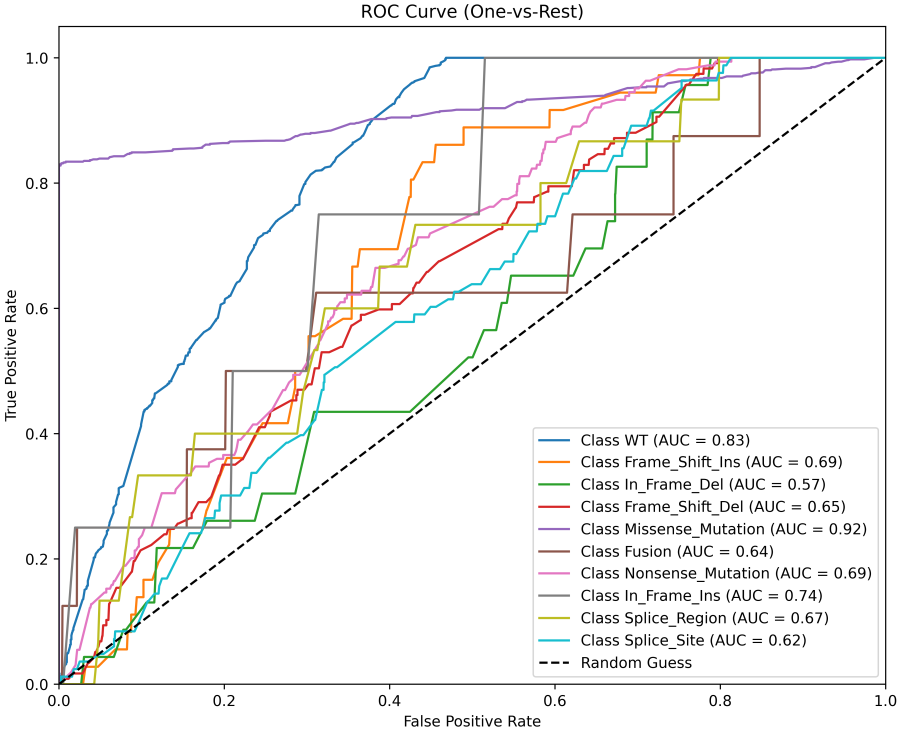


Figure S4: ROC curve for our Graph-Neural network classifier across all different classes each time in CCLE (top) and TCGA (bottom). We can see that the classifier performs reasonably on prevalent mutation types, specifically the WT and missense samples in abundance, but struggles with rare types. The lower classification performance is attributed to the limited number of training samples for the less prevalent classes, even with stratified labels in the train-test split and class weights applied in the loss function to favor less the dominant classes.

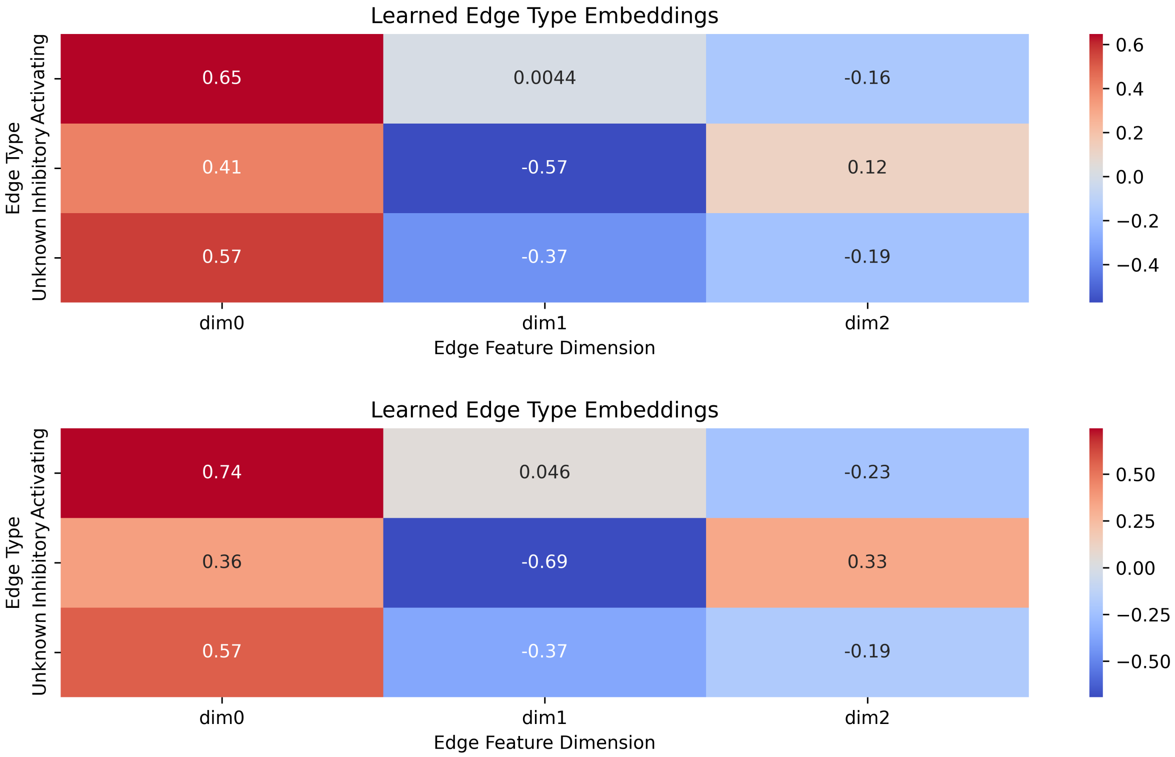


Figure S5: Learned edge attribute embeddings across CCLE (top) and TCGA (bottom). The one hot encoding of the mode of regulation was captured by different weights, indicating its significant role in the classification.

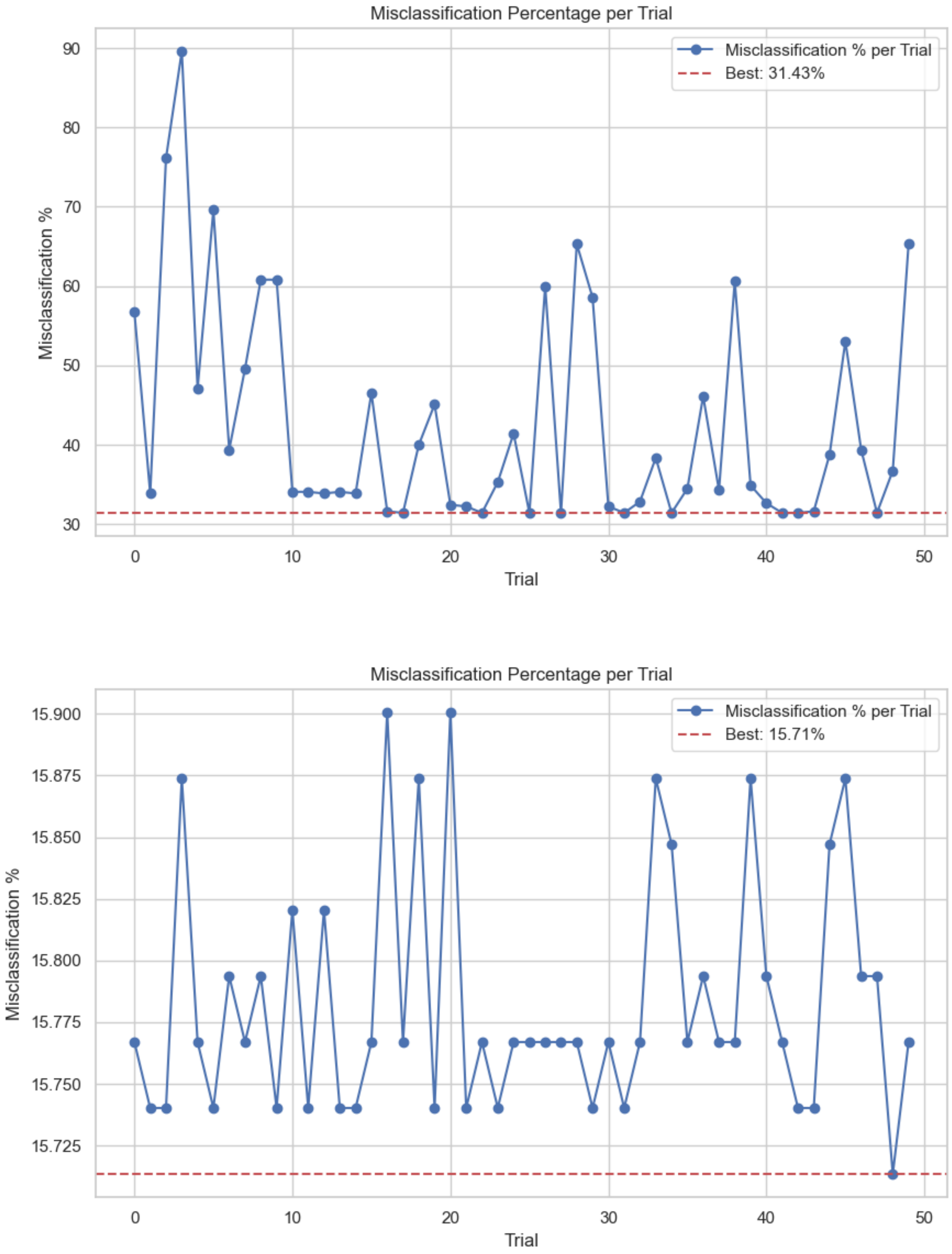


Figure S6: Optuna hyper-parameter tuning results for CCLE (top) and TCGA (bottom).
